# P4ward: an automated modelling platform for Protac ternary complexes

**DOI:** 10.1101/2024.11.28.625921

**Authors:** Paula Jofily, Subha Kalyaanamoorthy

## Abstract

Proteolysis Targeting Chimeras (Protacs) are a new class of drugs which promote degradation of a protein of interest (POI) by hijacking the Ubiquitin-Proteasome system. Structural knowledge of an E3 ligase: Protac:POI ternary complex is required for Protac rational design, and computational modelling of such heteromeric complex structures is nontrivial. To date, few programs have been developed to address this challenge, however, there remains a need for readily accessible tools that can significantly improve ternary complex modelling accuracy. Particularly, programs that can also support the screening phase of Protac discovery, where speed and the ability to test multiple Protacs is essential to advance the field of Protac therapeutics. To bridge these gaps, we present P4ward, a free and fully automated Protac ternary complex modelling pipeline. P4ward achieves a hit-rate of 76.5% with an average rank of 7.26, and substantially reduces the rank of the near-native pose by 73-98% compared to earlier programs. We believe that P4ward could be a user-friendly, fast, and effective tool for gaining atomistic insights necessary for Protac modelling and optimization.

## 1 Introduction

Proteolysis Targeting Chimeras (Protacs) are a novel class of therapeutics which promote the degradation of a Protein of Interest (POI) by hijacking the Ubiquitin Proteasom System (UPS), the endogenous cellular mechanism for regulated protein degradation. A Protac is a heterobifunctional molecule composed of three moieties: a POI binder (also called “warhead”), an E3 ubiquitin ligase binder, and a linker that connects these two ends. By binding to both proteins, the Protac induces the proximity between POI and E3 which would not otherwise occur, and consequently catalyses the POI’s destruction through ubiquitination. The interaction between these three components is key to successful degradation, culminating in the formation of a Ternary Complex (TC), where the Protac interacts with both proteins and, in many cases, initiates direct protein-protein interactions (PPIs). [1].

Since the establishment of targeted protein degradation (TPD) in 2001 [2], Protacs have emerged as a viable alternative to the conventional targeted protein inhibition methods [3, 4]. TPD offers multiple advantages over targeted protein inhibition, including (i) binding site versatility: unlike targeted protein inhibitors, which require orthosteric or allosteric binding sites on the POI, TPD only requires that the POI bind to the Protac’s warhead. (ii) cooperativity: TPD-enabled PPIs between the POI and E3 ligase can enhance the overall affinity of the TC when compared to the affinity of either binary complex (namely, Protac:POI or Protac:E3). (iii) degradation: TPD leads to to complete degradation of the POI, while the PROTAC molecule remains available for further cycles of therapeutic action. Consequently, Protacs act in substoichiometric concentrations, and even small doses can produce substantial, long-lasting effects [4, 1]. Given these benefits, Protacs are now promising candidates in field of the drug discovery, particularly for targeting proteins that are difficult to address with conventional methods [5, 6].

Recently, several Protacs have advanced in the clinical trials for the treatment of various cancers [5]. To facilitate rational design of Protacs, it is essential to understand the structural features of the ternary complex. Indeed, TC elucidation is an active area of research, and three-dimensional structures of ternary complexes have been used for guiding the rational design of potent Protacs [7, 8]. Currently, methods such as X-ray crystallography are used to obtain the three-dimensional structure of TCs, but lack the throughput necessary to support early discovery stages. Historically, computational tools have aided, accelerated and even enabled drug discovery. Computational structural biology can provide the framework for efficient Protac screening by accelerating linker design, TC modelling and assessment of degradation efficiency.

It is important to note that TC modelling bears additional layers of complexity when compared to conventional protein-ligand binary complexes due to the involvement of three-body interactions. Computational workflows must therefore be adapted, and new methods and pipelines must be developed specifically for Protac modelling. The pioneering study by Drummond *et al.* [9] tested four different approaches to modelling TCs, and showed that protein-protein docking provides the most viable starting point. Since then, there have been a few efforts in developing TC modelling tools. Rosetta developers employed their protein modelling tools with PatchDock and RDKit to develop PRosettaC [10], a webserver for single TC modelling. Drummond *et al.* later improved their best protocol [11], which now is available for use through the Molecular Operating Environment (MOE) software. Weng *et al.* [12] developed a Python2 script that integrates FRODOCK for PPI and RDKit for protac sampling. Bai *et al.* [13] used a similar approach by employing RosettaDock and OMEGA for PROTAC sampling. Rosetti *et. al.* [14] proposed a protac-centric workflow in MOE where linker conformational sampling, as opposed to protein-protein docking, is the main driver of TC structure sampling. ClusPro developers used a modified version of the protein-protein docking program PIPER and RDKit for PROTAC sampling to propose a modelling protocol[15]. Modelling pipelines specialized for Protacs are indispensable in their ability to consider all structure-driving components on the TC. Currently, there are no easily accessible platforms which perform large-scale, accurate, and automated TC complex modelling and screening. Here we present P4ward (Predictive Protacs Python Pipeline: workflow automation for research and development), a platform built to bridge the gaps in availabity, scalability and accuracy in the current scenario of Protacs TC modelling. P4ward is free and open-source, written in python3, and orchestrates protein-protein docking, linker sampling and scoring tools to generate TC structure predictions from simple user inputs. P4ward implements a comprehensive sampling and filtering protocol that considers full Cullin Ring models for accessible lysine check, protein-protac interaction scores, and internal quality assurance checks to promote a complete modelling experience. The program is also capable of sampling multiple linker variations in a single run, allowing users to perform virtual screening of multiple linker types for a warhead and E3 ligand pair. P4ward integrates permissively-licensed structural biology tools, is highly customizable and fully automated, and was developed with parallel processing support. P4ward is the first tool to provide easy and open access to fast and accurate prediction of Protac ternary complexes.

## 2 Results and discussion

### 2.1 Program overview

P4ward is developed as a python package, where each module is responsible for a main stage of the pipeline. Users interact with P4ward by providing a configuration file (.ini). A template configuration file with all possible parameters in their default values can be written by the program and edited as needed by the user. In the case where keywords are omitted by the user, the program will internally populate these keywords with default values. Final files from all modelling stages are written to the working directory, including a comprehensive table with all relevant results, interactive plots, and a visualization script for viewing final ternary complex predictions in ChimeraX [16]. Further, P4ward also enables advanced users to utilize its functions in their own scripts and tools by importing the package and its modules.

P4ward’s default workflow consists of four main stages of modelling: Molecular preparation, protein-protein docking, linker sampling, and protac scoring and clustering. These stages, as well as input and output, are outlined in Figure 1 and described in detail in Section 4.

**Figure 1:**
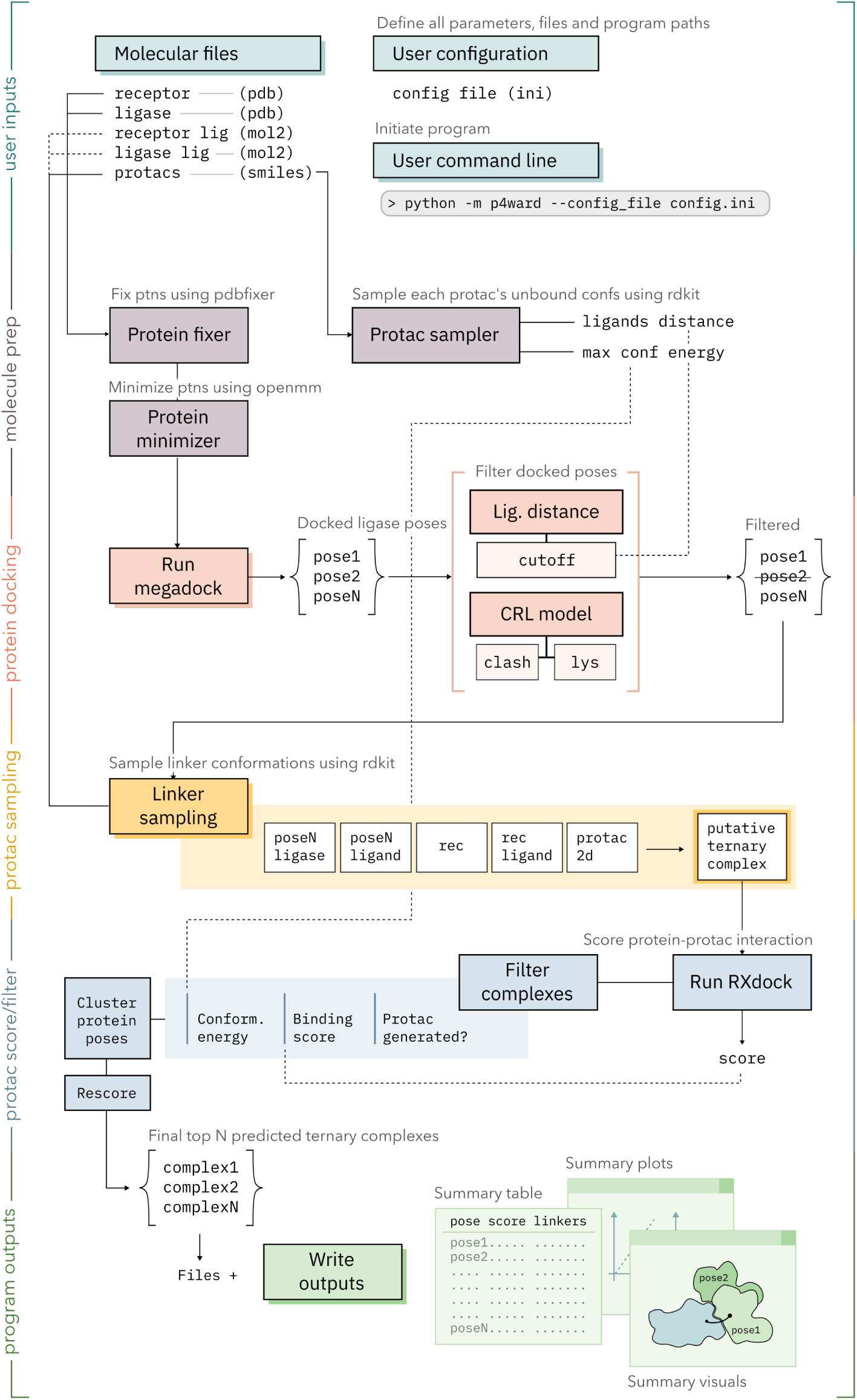
The default TC modelling workflow in P4ward. Different colors represent each stage of modelling. Boxes with darker shading represent steps which are not optional. Please see text, Section 4 for detailed description of each step. **Rec**: receptor, **Lig**: ligand, **Conf**: conformation. **CRL**: Cullin Ring Ubiquitin Ligase.

In brief, the proteins are prepared (fixed and minimized) using OpenMM and docked while bound to their respective ligands, using Megadock [17, 18]. Separately, unbound protac conformations are sampled to gather quantitative information about the molecule, which is used later to guide filtering steps. The protein poses are then filtered based on their binding site distances, as well as presence of accessible surface lysines for ubiquitination. At this point, low-cutoff clustering may also be performed to remove redundant poses. Next, Protac conformations are sampled onto the filtered protein poses using RDKit, while obeying the constraints of the bound ligands. Here, automatic checkpoints are applied to ensure structure sampling quality, please refer to sections 4.2 and 4.4 for more details. Next, the poses for which it was possible to sample protac poses are forwarded to rescoring, where protein-protac interaction energies are calculated using RXDock [19]. The newly generated ternary complexes are filtered to keep only favourable binding scores. At this step, a new, final score is generated to combine both protein-protein and protein-protac interaction scores, and the ternary complex models are then reranked based on this score. Finally, the models are clustered (referred to as trend clustering), and the cluster centroids are selected as representative structures, finalizing the P4ward results.

### 2.2 Validation

In order to test the ability of P4ward to accurately model ternary complexes, we built a benchmark dataset using the known TC structures available in the Protein Data Bank (PDB). The benchmark dataset included 36 reference TC structures, of which five are Cereblon-based (6BOY, 6BN7, 6BN8, 6BN9, 6BNB) and the remaining 31 are VHL-based protacs. These reference structures were subdivided into a training set, consisting of 14 diverse structures (12 VHL-based and 2-CRBN based), and a test set comprising the remaining data (Table S1). P4ward was benchmarked on this dataset through two main use-case scenarios: bound proteins and unbound proteins. The bound state scenario assumes the availability of an experimentally-determined TC structure, and P4ward is employed to evaluate the suitability of different linker compositions. The unbound state scenario replicates a use-case where models are being investigated for a receptor-ligase pair with no experimentally determined ternary complex. Because the proteins’ conformations are not already adapted to an interaction interface, this case is much more challenging to the pipeline than the bound case.

In the next sections, we describe P4ward’s configurations on the training set and validate its key modeling steps, including successfully sampling protac linkers based on predefined ligand positions, accurately identifying a productive protein pose in the context of the CRL model, and closely predicting native-like protac ternary complex structures. The best configurations identified from the training set are then used in the test set evaluations.

#### 2.2.1 Linker sampling

One of the key determining steps in successful modeling of TC is to sample linker conformations while respecting the spatial constraints of each protac’s ligands. Therefore, we assess P4ward’s RDKit-powered linker sampling capabilities by sampling linkers for the reference crystal structures. For each, we generate 10 linker conformations. Depending on the coordinates of the ligand atoms (receptor ligand and ligase ligand) and the strain they introduce, some conformations deviate from such coordinates and are considered unsuccessful and therefore discarded.

Conformers were successfully generated for 14/15 structures in the training set. The exception is the protac in 8QW6 (Table 1). We believe that the stretched linker in the crystal pose causes a high strain that cannot be reproduced by *in silico* sampling (Figure S1). Indeed, modelling of this TC structure in P4ward yields no valid conformations, leading to its exclusion from subsequent benchmarking studies. Notably, reference PDBs 7PI4 and 7JTP contain linkers comprising only one atom. In this case, P4ward was allowed to detect this and automatically extend flexibility into 2 neighbouring atoms (see section 4.4). All linker conformations for three PDB structures, 7PI4, 7JTP, and 8QVU, were discarded due to deviation of the ligand constraints (Table 1). However, this does not suggest a problem with the structures themselves and may reflect a limitation of the sampling procedure. Hence, we retained these structures in the final bench-marking dataset, which contains 14 PDB TC structures productive for modelling.

**Table 1:** Count of successful linker conformations sampled for each reference crystal structure.

| 7ZNT | 7S4E | 7PI4 | 7KHH | 7JTP | 6SIS | 6HR2 | 6HAY | 6HAX | 6BOY | 6BN7 | 5T35 | 8QW6 | 8QW7 | 8QVU |
| --- | --- | --- | --- | --- | --- | --- | --- | --- | --- | --- | --- | --- | --- | --- |
| 7 | 3 | 0 | 6 | 0 | 9 | 5 | 4 | 1 | 5 | 5 | 4 | - | 10 | 0 |

In addition, we test the accuracy of P4ward’s linker identification procedure. As described in detail in section 4.2, P4ward must identify, in the Protac 2D structure, the ligand atoms and the linker atoms. We show that when accurate structures are input, this step is 100% accurate (Figure S2). The linker matching check indicated that sanitization should be turned off for PDBID 7KHH, ensuring a perfect match for this structure.

#### 2.2.2 Accessible lysine filter

The accessible lysine filter considers a lysine accessible by Ubq if they are at most 60Å from each other, and if no other receptor protein atoms are found within a distance cutoff (lysine occlusion cutoff) of the line from Ubq C-terminal and the lysine’s side chain nitrogen (Figure 2).

**Figure 2:**
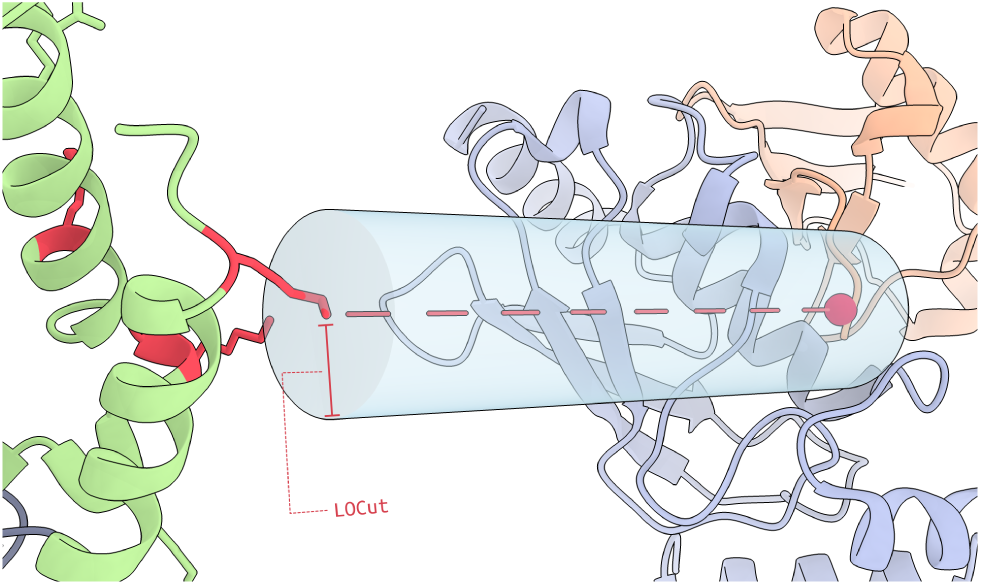
The Lysine occlusion cutoff. For a lysine to be considered accessible, no receptor protein atoms should be within this distance (represented by the blue cylinder) to the segment between the lysine and the Ubq C-terminal, represented by the pink dashed line. LOCut: lysine occlusion cutoff.

The lysine occlusion cutoff directly impacts how many lysines are considered accessible in each protein pose. In order to determine a default value for this cutoff and validate P4ward’s accessible lysine filter, we obtain the crystal pose for each of the ternary complexes in our bench-marking set, under the assumption that these poses would contain at least one accessible lysine. We then calculate their number of accessible lysines using increasingly stringent lysine occlusion cutoff values (Table S2).

As the lysine occlusion cutoff value increases, the number of accessible lysines decreases. At a cutoff of 4Å, no accessible lysine is identified for structure 7PI4; however, all others still have accessible lysines as it continues to increase until 6Å, where another structure no longer is identified. At 7Å, 6 poses do not have accessible lysines, which suggests this cutoff is too high. A higher cutoff where reference poses still have accessible lysines ensures that fewer protein poses proceed to the next sampling stage, and those are more likely to be higher-quality candidates. For this reason, and to minimize the influence of 7PI4 as an outlier, we selected 5Å as the default value for the lysine occlusion cutoff.

#### 2.2.3 Crystal pose reproduction

Following the selection of a threshold value for accessible lysine filter, we assessed P4ward’s ability to reproduce the 14 TC reference structures in both bound and unbound state scenarios. An advantageous feature available in P4ward is its ability for self-benchmarking. P4ward will run the modelling functions with no knowledge of the known crystal pose, then compare the poses in the final predictions with the known pose, as reference, using the Capri protein-protein interaction assessment [20, 21]. After benchmarking, P4ward will automatically add each model pose’s Capri prediction quality categories (high, medium, acceptable, incorrect) to the final result table.

##### 2.2.3.1 Training set evaluations

In order to determine the optimal set of parameters for achieving high success rates in TC modeling, we assessed various combinations of configurations, namely: (i) number of protein poses sampled by Megadock, (ii) toggling redundancy clustering and trend clustering, (iii) applying CRL model filtering, and (iv) toggling RXDock minimization. Training set calculations were performed on 40 Intel Skylake cores (2.4 GHz) or 40 Intel CascadeLake cores (2.5 GHz). We number-code these configuration combinations as 001-008 for the bound dataset and 101-109 for unbound dataset. Additionally, we assess the ranking of TC predictions based on P4ward’s final combined score or Megadock’s PPI score alone. We consider a modelling run to be a “hit” when at least one of the predicted TC conformations is of acceptable, medium or high-category protein pose as per the Capri assessment. Thus, for each configuration code, we report the number of hits (out of 14) achieved by P4ward, the average ranking at which the best-scoring hit was placed, as well as how many of these can be found at the top-10 ranked models (Table 2). It is important to note that, although the training set is used here to find the best configuration for P4ward, no structural features from this set are used in the training of the program. In other words, we refrain from utilizing any observable characteristics of the training data itself, such as PPI surface or presence of hydrophobic patches, for tweaking the program’s performance. This helps ensure that, in the face of a limited benchmarking set, we do not overfit to non-representative data.

**Table 2:**
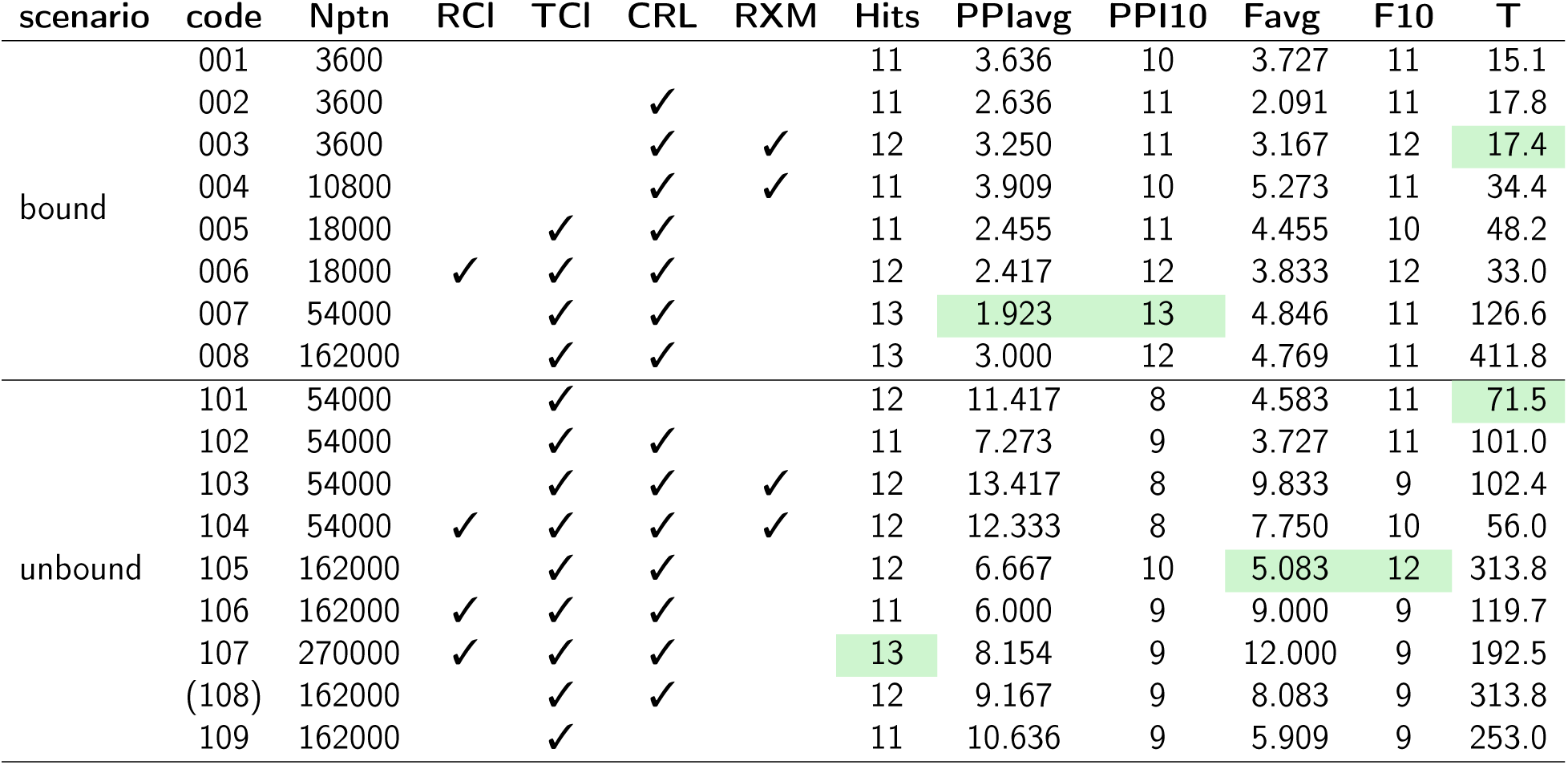
Description of the tested P4ward options for each configuration code. **Nptn**: number of protein docking poses generated by Megadock; **RCl**: toggled redundancy clustering; **TCl**: toggled trend clustering; **CRL**: toggle the CRL filter (includes clash and accessible lysine filter); **RXM**: toggle minimization of the protac bound to both proteins by RXDock; **T**: average run time, in minutes. **PPIH**: number of total hits, out of 14, when ranking by PPI score; **PPIavg**: average ranking by PPI score; **PPIdev**: Standard deviation of ranking by PPI score; **FH**: number of total hits, out of 14, when ranking by P4ward’s final score; **Favg**: average ranking by final score; **Fdev**: standard deviation of ranking by final score. Most accurate configurations for each scenario are highlighted in green. Config. 108 is identical to config. 105, except that Trend Clustering was done by selecting the highest scoring pose (based on the Final score) as the cluster representative, rather than the centroid, as used in all other configurations. Highlighted are the best results in terms of hit-rate, accuracy, or run time.

| scenario | code | Nptn | RCI | TCI | CRL | RXM | Hits | PPIavg | PPI10 | Favg | F10 | T |
| --- | --- | --- | --- | --- | --- | --- | --- | --- | --- | --- | --- | --- |
| bound | 001 | 3600 |  |  |  |  | 11 | 3.636 | 10 | 3.727 | 11 | 15.1 |
|  | 002 | 3600 |  |  | ✓ |  | 11 | 2.636 | 11 | 2.091 | 11 | 17.8 |
|  | 003 | 3600 |  |  | ✓ | ✓ | 12 | 3.250 | 11 | 3.167 | 12 | 17.4 |
|  | 004 | 10800 |  |  | ✓ | ✓ | 11 | 3.909 | 10 | 5.273 | 11 | 34.4 |
|  | 005 | 18000 |  | ✓ | ✓ |  | 11 | 2.455 | 11 | 4.455 | 10 | 48.2 |
|  | 006 | 18000 | ✓ | ✓ | ✓ |  | 12 | 2.417 | 12 | 3.833 | 12 | 33.0 |
|  | 007 | 54000 |  | ✓ | ✓ |  | 13 | 1.923 | 13 | 4.846 | 11 | 126.6 |
|  | 008 | 162000 |  | ✓ | ✓ |  | 13 | 3.000 | 12 | 4.769 | 11 | 411.8 |
| unbound | 101 | 54000 |  | ✓ |  |  | 12 | 11.417 | 8 | 4.583 | 11 | 71.5 |
|  | 102 | 54000 |  | ✓ | ✓ |  | 11 | 7.273 | 9 | 3.727 | 11 | 101.0 |
|  | 103 | 54000 |  | ✓ | ✓ | ✓ | 12 | 13.417 | 8 | 9.833 | 9 | 102.4 |
|  | 104 | 54000 | ✓ | ✓ | ✓ | ✓ | 12 | 12.333 | 8 | 7.750 | 10 | 56.0 |
|  | 105 | 162000 |  | ✓ | ✓ |  | 12 | 6.667 | 10 | 5.083 | 12 | 313.8 |
|  | 106 | 162000 | ✓ | ✓ | ✓ |  | 11 | 6.000 | 9 | 9.000 | 9 | 119.7 |
|  | 107 | 270000 | ✓ | ✓ | ✓ |  | 13 | 8.154 | 9 | 12.000 | 9 | 192.5 |
|  | (108) | 162000 |  | ✓ | ✓ |  | 12 | 9.167 | 9 | 8.083 | 9 | 313.8 |
|  | 109 | 162000 |  | ✓ |  |  | 11 | 10.636 | 9 | 5.909 | 9 | 253.0 |

P4ward achieves high accuracy (12/14 = 85.7%) in both bound and unbound scenarios (Table 2, Figure 3).

**Figure 3:**
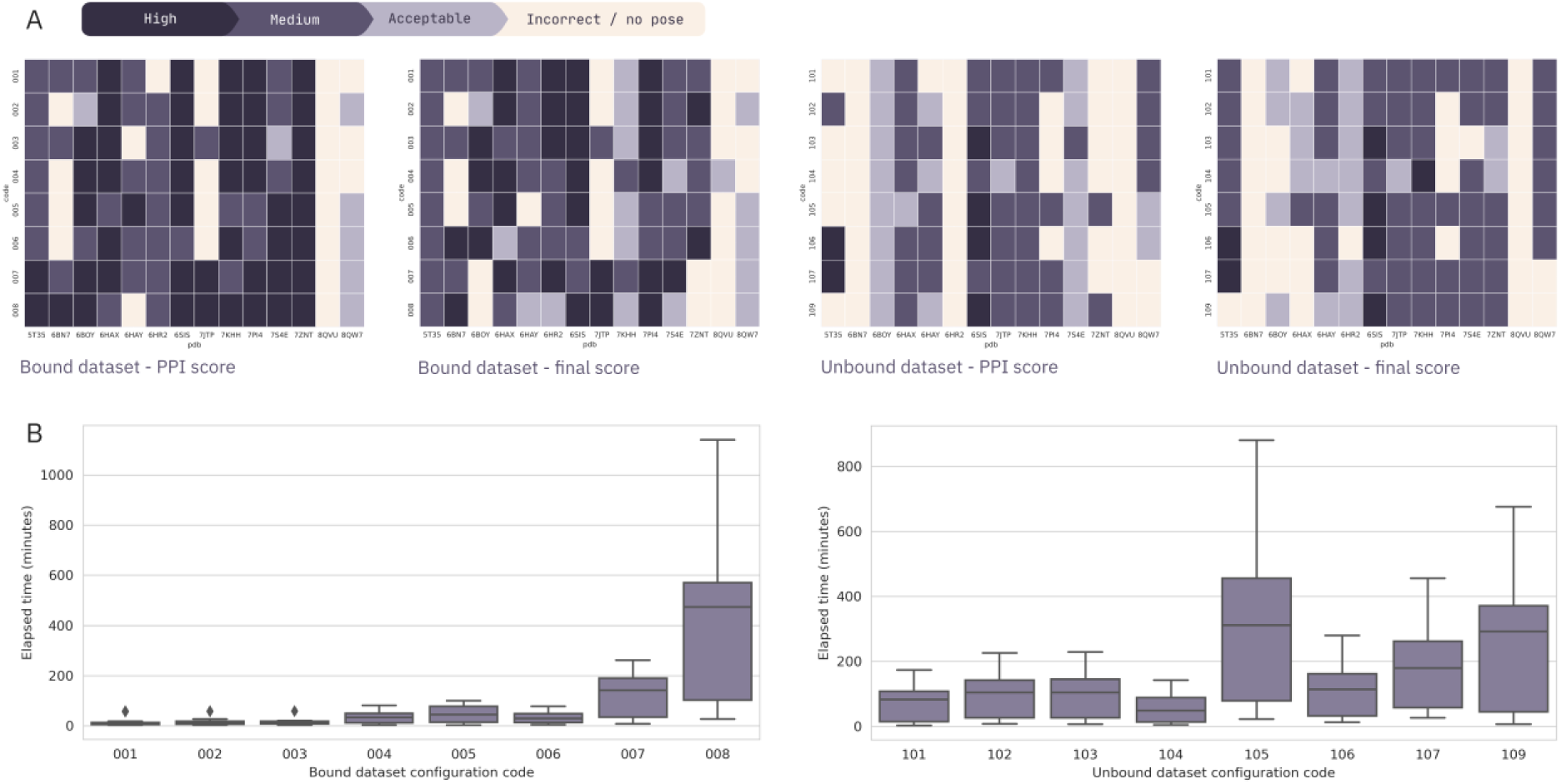
(A) Number and quality of hits for bound and unbound benchmarking scenarios on the training set, ranked by PPI or final score. For each configuration code and each reference TC structure, the color code describes the highest-quality category found at the corresponding top 10 ranking. (B) P4ward’s runtime for each configuration code in both bound and unbound scenarios.

###### Bound scenario

On the bound state scenario, fast and minimal sampling options (namely 3600 poses, PPI score, and no filters - config. 001) were able to achieve 11/14 hits with an average ranking of 3.25. This is due to the ease with which the protein interfaces can be identified by their shape complementarity and, for the same reason, it is advantageous in this case to rank by PPI score alone. A typical run with these settings would require less than 20 minutes, and could therefore be leveraged to screen a large number of new potential protac linkers for a known protac TC structure. We also show that with more exhaustive sampling and filtering, it is possible to achieve 13/14 hits with average ranking of 1.9 (config. 007), an increase of 18% on the hit-rate. It is also worth mentioning that there is a more pronounced increase in the quality of the generated poses. For example, at its top 10 ranked poses, config. 008 predicts 9 high-quality poses as opposed to 5 in config. 001. However, the use of config. 008 requires significantly more runtime – almost 7 hours – compared to 20 minutes for config. 001.

###### Unbound scenario

Currently, with very few known protac TC structures and the cost of experimentally solving protein structures, the unbound scenario represents the large majority of P4ward’s use case: predicting TC structures where none is known. Identifying protac-induced PPIs is challenging since such interactions are naturally unfavourable and happen only due to the protac’s catalysis. Additional challenges specific to TC in rigid-body docking include the limited surface area of the PPI and the poor shape complementarity of the binding regions in the unbound state. As a result, PROTAC-mediated PPIs are ranked lower by Megadock, which favours PPIs with better shape complementarity and larger surface areas. As expected, minimal filtering and PPI scoring result in an average rank of 11.417, with only 8 structures with top-10 hits, a significantly lower rank when compared to the bound dataset.

We also show that our implementation of the accessible lysine filter, trend clustering and the inclusion of protein-protac interaction scores significantly increases P4ward’s average ranking. For instance, combining protein-protac interaction scores with the PPI interaction scores increases the average rank to 4.583 on config. 101. Similarly, config. 107 achieves a higher hit-rate of 13/14 hits, although with a lower average ranking of 12.0. Notably, on config. 105, P4ward achieves a hit-rate of 12, where all 12 cases contain the best-scoring hit on the top-10 predicted TC models. The modelling steps implemented in P4ward efficiently filter out irrelevant PPIs and poses that do not support productive protac PPI formation, ensuring that ranking is performed on a refined pool of poses, reducing the risk of false positives. Additionally, trend clustering consolidates highly populated false-negative PPIs, minimizing their influence in the ranking to a single representative pose. This approach reserves top-ranked positions for low-ranking true positives that may otherwise be overlooked.

Further, P4ward’s final score significantly improved ranking across most configurations. Both the PPI score calculated by Megadock and the protac-protein interaction score by RXDock contribute equally to the final score. Since the protein-ligand interactions between the protac’s ligands and their respective proteins do not change across poses, the differences in protein-protac scoring arise from the linker conformations or a ligand’s interaction with the opposing protein. Therefore, assessing protein-ligand interactions allows for better TC modelling predictions, and accounts for the protac’s contribution to the TC’s cooperativity.

We have also found that redundancy clustering and RXDock minimization negatively impact ranking process. For example, the addition of redundancy clustering at config. 106 negates the significant improvement seen in the ranking with final score in config. 105; and the addition of RXDock minimization between config.s 102 to 103 drastically increased average ranking. The purpose of redundancy clustering is to decrease runtime by discarding highly similar protein poses early in the workflow. All protein poses which pass the distance filter are clustered on a 3Å distance cutoff and the centroid of the cluster is selected as the representative pose. If the cluster contains only two poses, the best scoring pose is selected. We reason that this step impacts the final ranking since it discards poses outside of the context of evaluating a Protac, which can be detrimental if potentially productive poses get discarded. Alternatively, RXDock minimization refines the protac’s conformation when bound to both receptor and ligase pose, allowing the linker coordinates to adjust after being initially sampled without the knowledge of the PPI. However, potent protacs include linkers that naturally promote productive PPIs. Therefore, productive PPIs (formed by true positive protein poses) enable sampling of linker conformations that do not require further adjustment. By skipping minimization, the poses that promote unfavourable linker conformations are discarded.

In light of this, we show that P4ward configured with CRL filter, trend clustering and Final score, and excluding RXdock minimization and redundancy clustering, yields the best accuracy and robustness for the unbound scenario (Table 2) – config. 105. In this configuration, P4ward achieves 12/14 hits, all favourably ranked. When analysing the top 10 models for each hit, 9 of the hits contain at least one medium-quality prediction, one features a high-quality structure, and only 2 of them contain structures of quality no better than acceptable (Figure 3). Figure 4A and Table S3 describe in detail the benchmarking results for config. 105, and Figure 4B shows that high- and medium-quality predictions reproduce protac conformations with high-accuracy.

**Figure 4:**
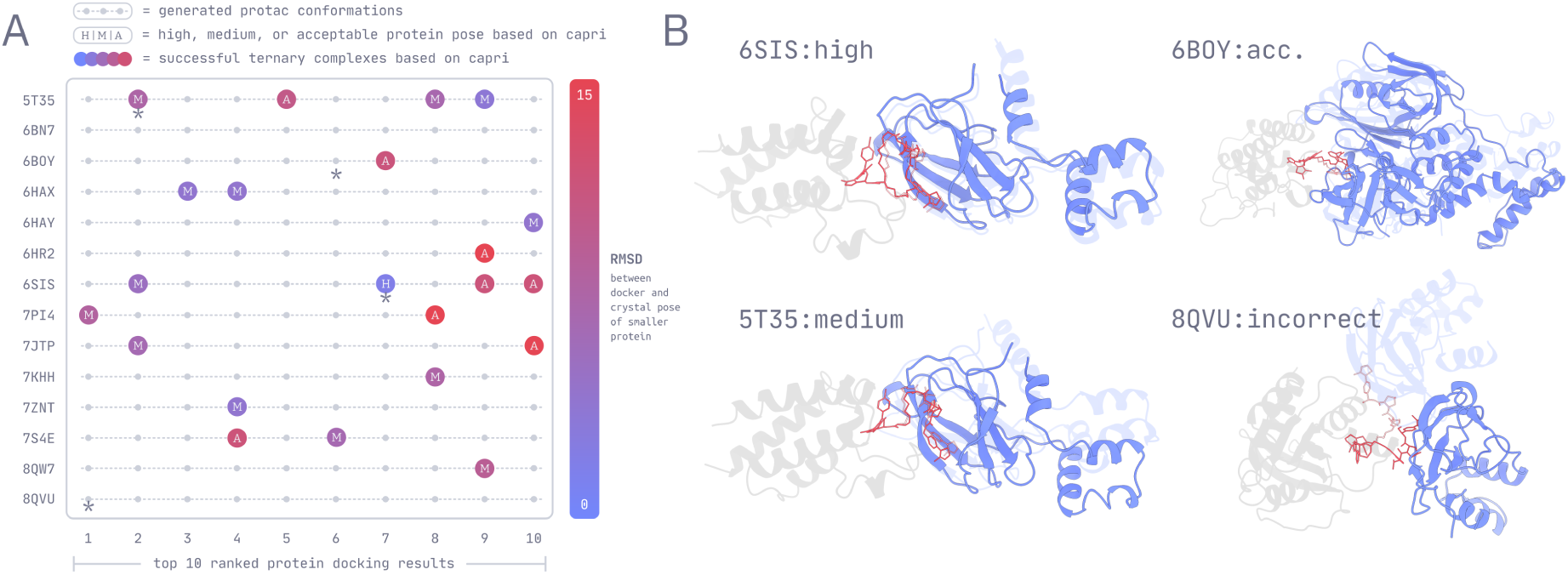
P4ward’s results in its best-ranking configuration (conf. 105). (A) Description of the top 10 predicted TC poses for each referece PDB. Colored spheres mark which poses were hits, with the embedded letter indicating which capri category the pose belongs to. Small gray spheres mark incorrect poses. Asterisks mark which poses are shown in (b). (B) Examples of modelled poses for each capri category. Light gray: receptor protein. Light blue: known ligase pose, Blue: Predicted ligase pose, transparent pink sticks: known protac binding mode. Solid pink sticks: predicted protac binding mode. See full CRL models in Figure S3.

We note that P4ward generates high-quality predictions for reference PDBID 6SIS in four of the configurations in the unbound scenario, and yields medium-quality predictions in all others (Figure 3). This is striking, given the strict Capri metrics which determine a protein pose to be of high-quality. Interestingly, 6SIS consists of a macrocyclic protac rationally designed to lock the TC onto the conformation previously identified and published as PDBID 5T35, thus yielding lower TC K_d_ [7]. P4ward is able to reflect this phenomenon: fewer protein poses can successfully model protacs in 6SIS if compared to 5T35, and those are concentrated into fewer trend clusters and exhibit higher enrichment of quality poses.

It has been shown that crystallographic structures such as the ones employed in this bench-marking study are but one of many possible low-energy conformations adopted by the TC [22, 23]. Experimental conditions such as crystal packing may also induce TC conformations distinct from a global minimum in solution [24]. In light of this, it is possible that other top-ranked predictions by P4ward consist of favourable low energy conformations not captured by the crystal studies. Recently, a comprehensive study by Dixon *et al.* [22] employed hamiltonian replica exchange molecular dynamics, principal component analysis and clustering to achieve a set of three most representative conformations for TCs of SMARCA2-Protac2-VHL (PDBID 6HAX), SMARCA2-Protac1-VHL (PDBID 6HAY), and SMARCA2-ACBI1-VHL (PDBID 7S4E). Notably, they show that the lowest free energy structures found for these systems may be distinct from the crystal pose. Protac2 in particular demonstrated a more dynamic free energy landscape, and its most representative cluster lies 13.5Å RMSD away from the crystal pose. P4ward was able to predict a TC conformation similar to this cluster (5.968Å lRMSD, Table S4). This pose was ranked 10th in config. 105 and, in the benchmarking studies, was considered incorrect by the Capri assessment (Table S3), suggesting that P4ward is able to predict relevant TC poses which are not captured by crystallographic studies.

We then re-evaluated P4ward’s top 10 predictions for each of the Protac systems studied by Dixon *et al.* [22] with their three predicted clusters as reference poses. We show that it was able to generate top 10-ranked hits for 6 of all 9 evaluations: all except 6HAY-Cluster0, 6HAY-Cluster2, and 7S4E-Cluster1 (Table S4). Due to similarities between some clusters and the crystal poses, many poses which are hits against the cluster-references are also hits against the crystal-references, however, in 4 of the 9 evaluations, a cluster-hit had been ranked “incorrect” against the crystal reference in the original benchmarking evaluation. Therefore, we suggest that in the unbound scenario P4ward can identify productive TC conformations beyond what has been determined in crystal studies.

##### 2.2.3.2 Test set evaluations

We selected the best P4ward configurations identified from the training set: codes 003 and 007 for the bound scenario and codes 101, 105, and 107 for the unbound scenario. Codes 003 and 101 presented a very short run time while retaining a high hit-rate and low average ranking, while codes 007 and 105 presented highest accuracy. Although with significantly lower average rank, code 107 was also selected due to its high hit-rate. We then evaluate these configurations for the 20 reference PDB structures which comprise the test set (Table S1).

The ligand match check revealed improper matches for PDBIDs 8PC2 and 6BN9, thus sanitization was turned off for these structures. Next, due to the structural complexity of their Protacs, codes 6BN8, 8BB2, 8FY0, 8FY1, 8FY2, and 6ZHC required increased sampling of Protac conformations, which were then increased to 100 from the default value of 10.

Seven of the structures included in the test set exhibit very minimal PPI, which challenges P4ward’s accuracy, since the program is aimed at TC’s with PPIs and depends on protein docking as an initial model generator. Additionally, reference structure 6BN8 exhibits a 10 PEG units-long linker, which results in a distance filter cutoff of 33.87Å, challenging P4ward’s accuracy by not significantly reducing the number of poses sent to the subsequent modelling stages. Further, since 6BN8 includes the more flexible CRL4^CRBN^ E3 ligase, fewer poses are also discarded by the CRL filter. For this reason, the test set was subdivided into a challenge set, comprising these eight reference poses, and a regular set, comprising the remaining 12 reference structures. The Capri benchmark assessment takes into account three parameters: RMSD of the smallest protein, interface RMSD, and fraction of native contacts (FNat), which is determined as the number of correct residue-residue contacts divided by the number of contacts in the target (or reference) complex. The challenge set comprises of structures with minimal interface and thus very few contacts in the reference structure. This generated very large values for FNat when benchmarking these structures, which led to the miscategorization of incorrect poses as acceptable or even medium-quality poses. To ensure our benchmark does not include false-positives, we re-calculated the challenge set results, excluding the FNat contribution. Table 3 reports the number of hits and ranking statistics for PPI score and Final score for the chosen configuration codes on the test set.

**Table 3:** Description of P4ward’s results on the test set, using the best performing configurations identified on the training set. **PPIH**: number of total hits when ranking by PPI score; **PPIavg**: average ranking by PPI score; **PPIdev**: Standard deviation of ranking by PPI score; **FH**: number of total hits, when ranking by P4ward’s final score; **Favg**: average ranking by final score; **Fdev**: standard deviation of ranking by final score. Benchmarking of the challenge set did not take into account all Capri parameters, please see text for description. Values marked with (*) are from a single datapoint.

| Scenario | code | Regular set (12 structures) |  |  |  |  | Challenge set (8 structures) |  |  |  |  |
| --- | --- | --- | --- | --- | --- | --- | --- | --- | --- | --- | --- |
|  |  | hits | PPlavg | PPI10 | Favg | F10 | hits | PPlavg | PPI10 | Favg | F10 |
| Bound | 003 | 8 | 3.125 | 8 | 5.75 | 7 | 3 | 9.333 | 2 | 8.667 | 2 |
|  | 007 | 9 | 6.889 | 7 | 9.444 | 7 | 1 | 4.0* | 1 | 1.0* | 1 |
| Unbound | 101 | 8 | 9.375 | 5 | 14.0 | 6 | 4 | 9.5 | 2 | 2.25 | 4 |
|  | 105 | 9 | 5.667 | 7 | 8.111 | 6 | 5 | 13.4 | 2 | 8.6 | 3 |
|  | 107 | 9 | 7.222 | 7 | 7.889 | 7 | 4 | 17.0 | 2 | 5.25 | 4 |

As expected of the performance of a test set when compared to the training set, the number of hits of each configuration decreases overall. P4ward can achieve 9/12 (75%) in the regular test set for both bound and unbound scenarios. Excitingly, it is also able to produce up to 5/8 (62.5%) hits in the challenge test set. In this case, where the PPIs were limited, the average ranking of the hits was improved significantly by the final score, emphasizing the role of protein-protac interactions in Protac TCs. On the other hand, PPI ranking performed better on the regular set, where it is likely that PPIs drive cooperativity to a greater extent than protein-protac interactions. In this case, a higher average rank for final score resulted in fewer top 10-hits for most configurations.

On all configurations except 003, the large cutoffs in reference structure 6BN8 proved computationally intractable. The calculations were not completed and 6BN8 was considered a non-hit in the reported results.

###### Bound scenario

In the bound scenario, we observe that config. 003 performs best overall, with better ranking on the regular set and three times as many hits on the challenge set (we do not consider the average ranks for config. 007 significant on the challenge set since there was only one hit, thus they arise from a single data point). We conclude that trend clustering in the bound scenario (007) can improve hit rate across the training and regular test sets, however, in the challenge set, both configurations would require higher protein pose sampling to improve accuracy. There is also no pronounced difference on the number of medium and high-quality poses between configs. 003 and 007 (Figure 5). Therefore, due to its high accuracy in the context of very low computational time, we consider config. 003 P4ward’s best candidate for bound TC modelling studies.

**Figure 5:**
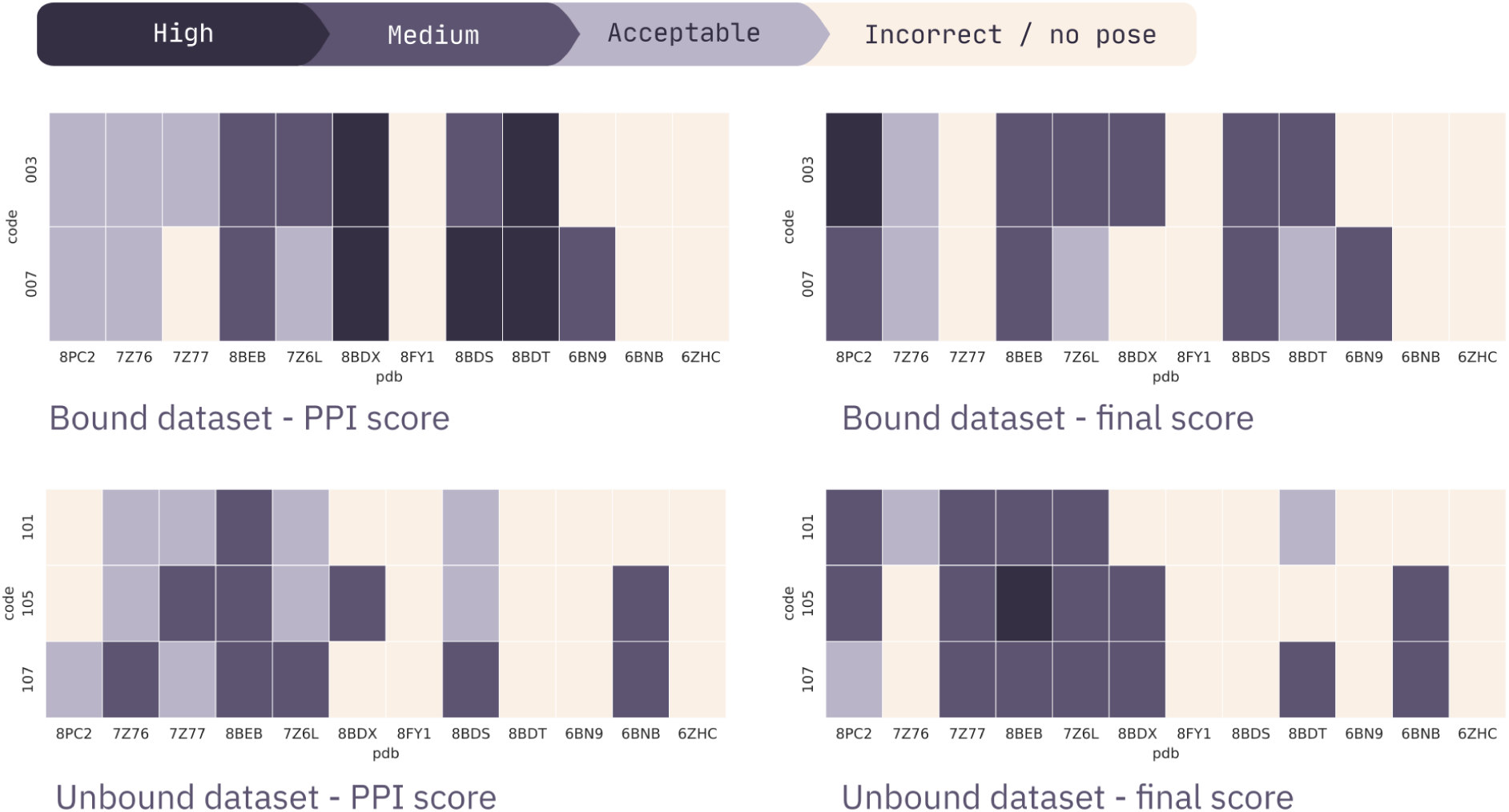
Number and quality of hits for bound and unbound benchmarking scenarios on the test set, ranked by PPI or final score. For each configuration code and each reference TC structure, the color code describes the highest-quality category found at the corresponding top 10 ranking.

###### Unbound scenario

In the unbound scenario, config. 105 achieves the best hit-rate in both regular and challenge sets, low average rank on the regular set, as well as higher-quality poses on the top 10 predictions (Figure 5, Table S5). Despite exhibiting the lowest top 10-hits for the challenge set, its average ranking for Final score is below 10, and indeed no hit was ranked lower than 15^th^. Therefore, we designate config. 105 with Final score as the representative of P4ward’s accuracy and performance, and all default values in the pipeline were adjusted to reflect this configuration. Across the whole benchmarking set, P4ward achieves a 76.5% hit-rate with an average rank of 7.26, where 80.76% of the hits (61.76% of total) are found on the top 10 ranked predicted models. Excluding the challenge set, P4ward achieves 80.8% hit-rate with an average rank of 6.6, where 85.71% of the hits (69.23% of total) are at the top 10 predictions.

Recently, Rovers and Schapira [25] performed a comprehensive benchmarking study comparing the accuracy of the Protac TC modelling programs currently available: PRosettaC, MOE, and ICM, as well as the state of the art protein-protein interaction modelling tools Haddock and Alphafold. They show that Alphafold and Haddock are not suited for TC modelling because these tools do not consider the Protac, and therefore fail to yield low RMSD structures. Their assessment was performed in the unbound scenario, and includes the average rank, in percentage, of the lowest RMSD pose (near-native pose) generated by each program, as well as the total number of poses and the average RMSD value of the near-native pose. In order to directly compare with P4ward’s results, we perform the same assessment with config. 105, ranked by Final score (Table 4). Table S6 describes the similarity between the benchmarking set utilized by the authors and the set utilized in the present study.

**Table 4:** Comparison between the benchmarking results reported by Rovers and Schapira [25] and P4ward’s results on our complete benchmarking set. The **Rank** values for the other programs was calculated from **Rank(%)** and the **Total** output poses reported by Rovers and Schapira. All values are averages across the benchmarking data.

| Program | RMSD(Å) | Rank | Rank(%) | Total |
| --- | --- | --- | --- | --- |
| Haddock | 17 | 5.61 | 17 | 33 |
| AlphaFold | 21 | 10.5 | 42 | 25 |
| ICM | 5 | 513.38 | 38 | 1351 |
| MOE | 11 | 1814.4 | 32 | 5670 |
| PRosettaC | 13 | 95.37 | 51 | 187 |
| P4ward | 10.98 | 25.438 | 49 | 52 |

P4ward produces near-native poses with average RMSD comparable to PRosettaC and MOE, but ranked significantly lower if compared to any of the Protac TC modelling tools, suggesting that currently, P4ward exhibits the best ranking power when compared to available Protac TC modelling tools, and brings a 73% improvement in ranking power when compared to the best performing program, PRosettaC. This is because P4ward achieves an enriched, smaller set of final TC predictions. Thus, although P4ward’s rank percentage is no smaller than its counterparts’, the higher enrichment of total results ensures a much smaller absolute rank. We consider the absolute final rank to be highly important. In a modelling workflow, a program is employed to generate predictions and a limited-sized subset of the top models is selected to subsequent analysis stages. In the case of ProsettaC, ICM and MOE, this subset would have to consist of 95, 513, and even more than a thousand models, respectively, to include the best near-native pose. However, it is more feasible to include the top 25, as with P4ward. In addition, as previously discussed, 70-85% of P4ward predictions include at least an acceptable pose on the top 10. In light of this, we consider P4ward the first tool to bring powerful-enough ranking to satisfy the needs of a computational Protac discovery workflow. We conclude, therefore, that P4ward presents a valuable addition to the available computational methods for Protac TC modelling.

## 3 Conclusion

We have developed a freely available, accurate Protac modelling tool. P4ward consists on an automated pipeline which integrates structural biology and chemistry tools for protein-protein docking, protein structure manipulation and protein property calculations, protein-ligand docking, and small molecule modelling. P4ward also implements extensive code for building a modelling protocol specifically tailored to Protac ternary complexes. P4ward offers a unified platform to bridge the gaps in current Protac ternary complex modelling methods. Compared to previous TC modelling tools, P4ward delivers unique advantages. First, it is free, easy to install, and designed to lower access barriers, providing seamless usability across Unix-based platforms, including computational clusters. Second, it enables virtual screening of Protacs, by accepting multiple Protacs in a single input. Third, it provides an extensive set of configurable filters, such as accessible surface lysines, which can be used to customize a modelling run or to complement protac related studies. Finally, P4ward offers a substantial improvement on benchmarking accuracy on experimental TC structures, while also capturing other relevant conformations beyond the single local minimum represented by such structures. P4ward was benchmarked against known Protac ternary complex structures, achieving 76.5% hit-rate with an average rank of 7.26, where 80.76% of hits are found on the top 10 predicted models. We believe this program will contribute in accelerating Protac discovery and advancing low-cost access to Protac research. P4ward is available on Github at https://github.com/SKTeamLab/P4ward, and can also be obtained through Docker.

## 4 Methods

### 4.1 Program inputs

The user input files are: target protein (.pdb format) (Receptor, Rec) and its bound ligand (.mol2 format) (RecL), E3 Ligase Substrate Receptor (.pdb format) (E3) and its bound ligand (.mol2 format) (E3L) and a .smiles file containing a list of all protacs to be sampled. The pipeline supports screening multiple linkers as long as the ligands (E3L and RecL) remain the same; therefore, we refer to each of the protac smiles codes in the input file as Protac Linker Variant (PLV) (Figure 6).

**Figure 6:**
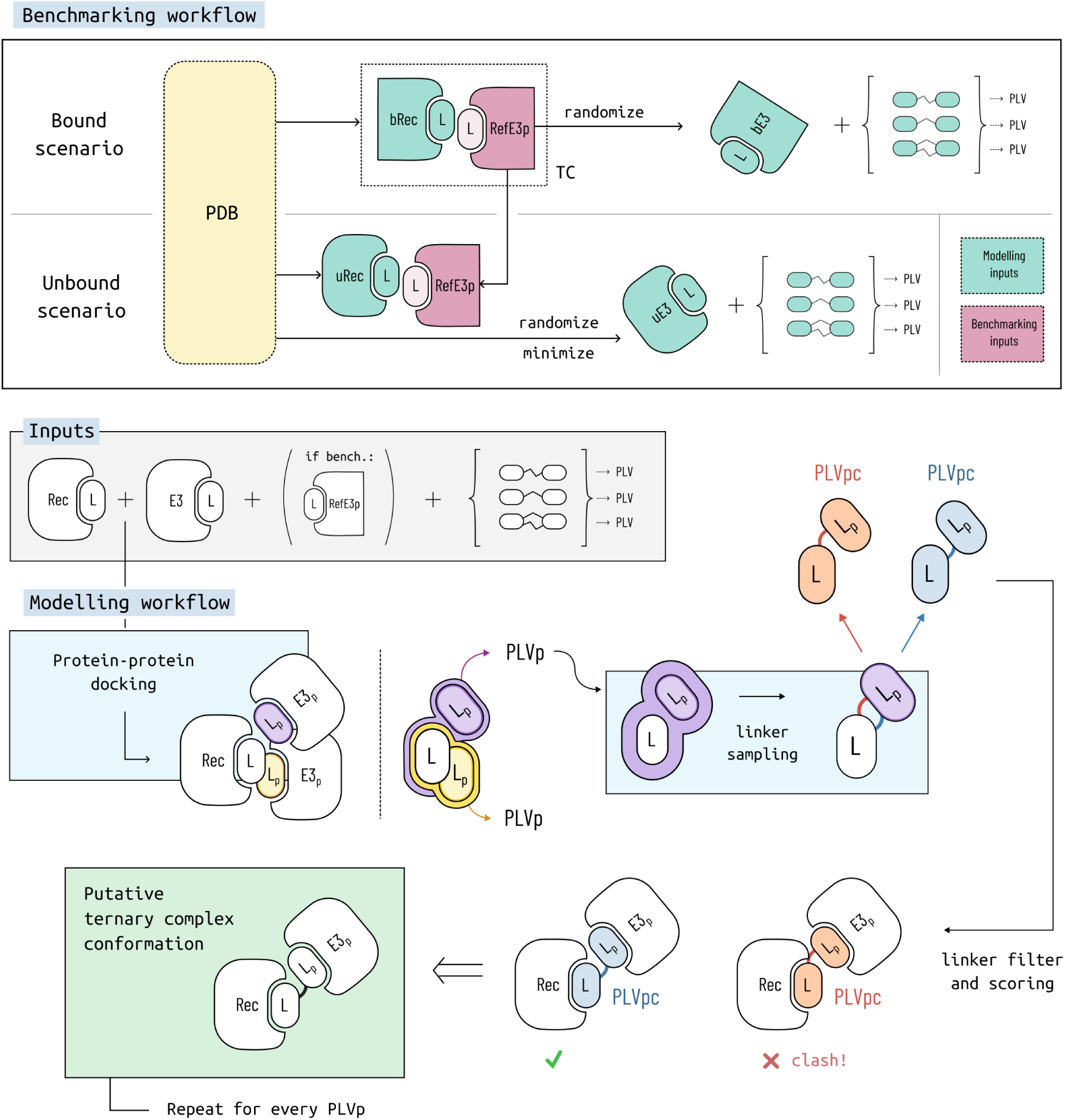
The structural and molecular components in the benchmarking and modelling workflow. **PDB**: Protein Data Bank;: Receptor - protein of interest; **E3**: E3 ligase used for docking onto receptor; **RecL**: ligand bound to the receptor; **E3L**: ligand bound to E3; **bRec/bE3**: Receptor and E3 in bound conformation, as extracted from the protac ternary complex structure; **uRec/uE3**: Receptor and E3 in unbound conformation, extracted from independent PDB entries; bRec, uRec, bE3 and uE3’s positions (with their ligands) were randomized for benchmarking. uRec and uE3 were minimized around their respective ligands; **RefE3_p_**: known binding mode of E3 onto Rec extracted from the ternary complex structure; **E3_p_**: a binding pose of E3 onto Rec generated by the protein-protein docking stage; **PLV**: protac linker variation - the complete 2D protac structure (smiles format); **PLV_p_**: the pose (excluding the linker) a protac would take when bound to Rec and a specific E3_p_; **PLV_pc_**: the possible linker conformations sampled for a PLV_p_ - includes the 3D conformation for the whole protac.

### 4.2 Molecule preparation

The first step in the P4ward pipeline involves preparation of the user input files (POI, E3 ligase). In the preparation step, P4ward uses the pdbfixer tool to add missing atoms and hydrogens, and can perform energy minimization the protein structures using the L-BFGS optimizer [26] in OpenMM.

P4ward leverages RDKit to interpret and sample the Protac molecule as well as the bound ligands in many stages of the pipeline. The user inputs RecL, E3L and the full protac 2D structure (as smiles code), without any explicit identification of linker atoms. P4ward automatically matches the ligands to the protac 2D structure and marks the ligand and linker atoms within the Protac molecule. P4ward can also generate an image file of the Protac structure, identifying the atoms as ligand and linker for user review prior to running the next step in the modeling pipeline.

In addition, RDKit is also used to sample each PLV’s unbound conformations. Thus, conformations are generated for each Protac molecule without any constraint introduced to the coordinates of the ligands that bind Rec and E3. These sampled conformations are used only to extract quantitative information about the Protac such as average linker length, and the conformations are not considered in the later TC modelling stages. This sampling is enabled when the user selects the usage of an automatic cutoff that calculates the proximity between the two binding sites (POI and E3, explained further in section 4.3). Large molecules with many rotatable bonds pose a challenge for achieving convergence in conformational sampling, often leading to the generation of only a subset of the number of desired conformations. For such cases, P4ward is designed to provide suggestions on increasing the number of requested samples for both unbound and bound conformations for linker sampling, discussed in section 4.4.

### 4.3 Protein-protein docking and protein pose filtering

P4ward implements protein-protein docking of the receptor and E3 (or their fixed/minimized versions) through Megadock, with Receptor being considered as static and E3 as ligand i.e., allowed to move during the docking calculation. In this step, a temporary .pdb file for each protein is generated to include its ligand respective. This file is used for docking, which allows Megadock to be aware of the ligands while docking the proteins. The pipeline parses the Megadock output file and captures translation and rotation information for each E3 docked pose (E3_p_). Using the rotation and translation vectors of E3_p_’s, P4ward transforms E3L so that it follows each new protein pose, thus generating a E3 Ligand pose (E3L_p_) for each E3_p_ (Figure 6). Following the pose generation, the distance between the centre of mass of the two ligands (RecL and E3L_p_) is calculated. If the distance is below the previously defined cutoff (see section 4.2, note that the cutoff value can also be set manually), the corresponding E3_p_ is forwarded to the next modelling stage of the pipeline.

#### 4.3.1 Cullin-RING E3 model

In addition to the classical protein-protein docking workflow, P4ward implements a dedicated filter for screening each E3_p_ in the context of the full Cullin-RING E3 ligase (CRL) ubiquitination complex: the E3_p_ is aligned to the full CRL model and filtered based on clashes with any of the CRL components, as well as availability of accessible lysine residues for ubiquitination. We have constructed CRL models CRL2^VHL^ and CRL4^CRBN^, which are included in the program for the accessible lysine check. Although the user may supply any E3 ligase substrate receptor as E3, currently the full model CRL filter can only be applied for VHL or CRBN. See supplementary section 1.1 for detailed modelling description.

P4ward aligns each E3_p_ that passed the distance filter to the CRL model. The resulting alignment matrix is then used to transform Rec, generating its pose bound to E3_p_ in the context of CRL (Rec_pCRL_). Next, the position of each Rec_pCRL_ within the complex is assessed by checking for clashes with any other CRL component. P4ward discards any E3_p_ that causes a clash and, if no clashes are found, checks for accessible lysines on the surface of the Rec_pCRL_. A lysine residue is considered potentially accessible if it is not occluded by Rec_pCRL_ atoms, has a solvent accessible surface area (SASA) greater than 2.5Å, and is located within 60Å [27, 22, 15] of the Ubq C-terminal glycine residue for CRL2^VHL^ or 16Å for CRL4^CRBN^. (more details on lysine occlusion in section 2.2.2). P4ward records the number of accessible lysines for each E3_p_. All E3_p_s that have at least one accessible lysine are selected for subsequent analyses. In the case of CRBN, where multiple CRL4^CRBN^ models exist, this process is repeated for each available model. The E3_p_ passes the filter if at least one model has an accessible lysine. All the cutoffs described in this section are default values in P4ward’s settings and can be modified by the user.

### 4.4 Linker sampling

RDKit is employed to sample conformations for each PLV based on the top N ranked E3_p_s by Megadock score (note that this is unrelated to the unbound conformational sampling mentioned in section 4.2). This step generates PLV conformations in the context of a putative TC (Figure 6). For a given RecL-E3L_p_ pair, a PLV adopts a putative pose, PLV_p_, where its ligand coordinates align with those of RecL and E3L_p_’s coordinates. For each PLV_p_, a configurable number of linker conformations (PLV_pc_) is sampled. Note that, if the linker is extremely small, such as one atom’s length, it is virtually impossible to generate a conformation for the PLV while keeping all other atoms rigid, because a small variation on the protein pose would render the linker’s structure implausible. In these situations, the pipeline has the ability to extend flexibility beyond the linker atoms into their closest neighbouring ligand atoms. Any PLV_pc_ that does not maintain the original ligand coordinates from PLV_p_is discarded. Subsequently, P4ward will analyse all PLV_pc_s: RDKit’s reported conformational energy is captured and RXDock is used to perform energy minimization of the PLV_pc_ in the context of Rec-E3_p_ and extract its binding energy.

#### 4.4.1 Protac scoring and filtering

Using P4ward’s simplest configuration, the top N-E3_p_s ranked by Megadock score are forwarded to protac sampling described in the previous section (4.4). These become the final results, independently of whether some of these cannot form viable protac conformations. Alternatively, the user may choose to extend sampling using the top ranked protein poses until N of those can successfully form protac conformations. In this case, P4ward will keep sampling PLV_p_s until N PLV_p_s have produced at least one successful PLV_pc_. By default, P4ward considers a PLV_pc_ successful if: (a) it can be sampled by RDKit, and (b) if its binding energy to Rec-E3_p_ is negative; the user may also choose to account for (c) if its conformational energy is lower than the maximum PLV unbound conformational energy. The user may apply filter combinations (a), (ab - default), (ac), or (abc). By applying these filters, the final results will contain N ternary complex conformations from the top ranked protein poses for which it was possible to generate a protac conformation with favourable binding energies (Figure 1).

Each putative TC conformation, which includes Rec, E3_p_ and its sampled PLV_pc_s, is rescored and reranked. The final score is generated by averaging the RXdock interaction score among all successful PLV_pc_s to achieve an array with a single RXDock protein-protac score and a single Megadock protein-protein score for each TC conformation. These are then l2-normalized and averaged to achieve a final score that is used for reranking all filtered TC conformations.

#### 4.4.2 Clustering

Clustering of protein poses can be carried out at two separate stages: after distance filtering, to remove very similar protein poses and speed up subsequent calculations (redundancy clustering, default cutoff is 3Å); and after protac sampling, to capture the trends in protein docking results (trend clustering). In the latter case, all the successful protein poses for each PLV are sent to clustering analysis (default cutoff of 10.0Å is used). For each protein pose, P4ward reports its cluster number, and whether it is the centroid and/or the best final scoring pose in that cluster. The clustering is performed using a simplified three-dimensional representation of the E3_p_s. Each E3_p_’s orientation in space is represented by three points, where the first point represents the E3L_p_ COM, the second point is in the segment between E3L_p_ COM and E3_p_ COM, 5Å away from E3L_p_ COM, and the third point is any point 5Å from the second point, perpendicular to the E3L_p_ COM-E3_p_ COM segment. This three-point representation allows for fast and accurate clustering with low RAM requirements. The clustering calculations are performed with the Hierarchical Agglomerative algorithm [28] with Ward linkage [29].

### 4.5 Outputs

Files generated from the main modelling stages are saved to the working directory: the fixed and minimized structures of Rec and E3, the relevant E3 docked poses, the sampled protac conformations for each filtered E3_p_, and the protac poses as minimized and scored with RXDock. P4ward also generates a summary folder for each PLV tested, with a table describing the ranking of all final TC models, and any combination of the following analysis: Data plots, 3D visualization scripts, and complete CRL complex models.

The plots generated by P4ward are interactive and written to a single HTML file which can be viewed in any web browser. Currently, P4ward generates a funnel plot showing how many protein poses were selected by each modelling stage; a protein-protein interaction distribution; 2D plot of Components 1 and 2 extracted from Principal Component Analysis of the protein poses’ 3D coordinates, coloured based on each passed filter; and a scatter plot of the final models’ PPI scores, Protein-Protac interaction scores, and combined final score.

Visualization scripts are written for the program ChimeraX and allow a complete view of all final predicted TC models. If benchmarking was performed, the reference ligase pose is also shown, and the protein poses are coloured based on their Capri rank.

Finally, if the user so chooses, P4ward can generate PDB files of the full CRL complex models for the final predicted ternary complexes.

### 4.6 Benchmarking preparation

In the bound state scenario structures, the Rec and E3 are in the exact same conformation as found in the TC, and we extract the proteins from the reference ternary complex PDB structure: the receptor protein (bound Rec structure, bRec) and the E3 protein. This E3 protein pose is the known experimental binding pose, therefore referred to as reference E3 pose (RefE3_p_). To remove any influence the initial position could have on Megadock’s results, we duplicate RefE3_p_ and randomly translate and rotate it in space, retaining the ligand (E3L). This step generates the bound E3 (bE3) and bound E3 ligand (bE3L). The TC PDB entry is also used to obtain the .mol2 file and smiles structure of the Protac molecule, and the bound receptor ligand (bRecL) and the bound reference E3 ligand (RefE3L_p_) are extracted (Figure 6).

In the unbound scenario, unbound Rec and E3 structures (uRec and uE3) are obtained separately from different PDB entries (Table S1 describes the PDBIDs used for the proteins of each reference TC). The uE3 structures are also randomly translated and rotated, and their respective ligands are aligned to their positions in space, generating uRecL and uE3L. This alignment can cause mild steric clashes, especially since not all unbound proteins are available as binary complexes with the full corresponding ligand. To remove these clashes but maintain the known binary binding poses, we further prepared the complexes by performing energy minimization of uRec and uE3 while restraining their ligands.

## Acknowledgements

This research was enabled by the countless contributors to open-source programs and packages as well as the high-performance computing provided by the Digital Research Alliance of Canada. We would also like to extend our thanks to our software testers, Max Walton-Raaby, Steven Ong, and Justine Williams.

## 5 Contributions

S.K. and P.J. conceived the project. P.J. developed the software. P.J. designed and conducted the experiment(s), analysed the results and drafted the paper with input from S.K. S.K. supervised the project and acquired funding. Both authors reviewed the paper.

## 1 Supplementary Text

### 1.1 CRL2^VHL^ and CRL4^CRBN^ models

For both CRL2^VHL^ and CRL4^CRBN^ models, PDBID 6TTU (containing Cul1 as scaffold protein and beta-TRCP as substrate receptor) was used as reference to obtain an active conformation of the CRL. We modelled the complex VHL-EloBEloC-Cul2-NEDD8-Rbx1-E2-Ubq from several PDB structures (referred to as S1 to S4) as follows: (S1) PDBID 6TTU was opened as reference. (S2) PDBID 5N4W provides structures for VHL (partial), EloC, EloB, Cul2, and Rbx1. We aligned S1 to S2 using residues 416 to 672 comprising the C-terminal domain of Cul2. Next, we obtained (S3) PDBID 1LQB which provides complete structures for VHL, EloC, EloB. We aligned S3 to S2 using EloCB residues, and delete VHL, EloC, EloB structures from S2. We then opened PDBID 5N4W again (S4) and extract NEDD8 (keep residues 688 to 759, delete all others). Lastly, S4 is aligned to the corresponding NEDD8 residues in S1.

For CRBN-DDB1-Cul4A-NEDD8-Rbx1-E2-Ubq, we followed the protocol described by Bai et. al. [27] as follows: (S1) PDBID 6TTU was opened as reference. (S2) PDBID 2HYE provides structures for Rbx1, Cul4A, and DDB1. We aligned S1 to S2 using residues 416 to 672 comprising the C-terminal domain of Cul4A. We then opened PDBID 2HYE again (S3) and extracted NEDD8 (keeping residues 688 to 759, and deleting all others). NEDD8 from S3 was aligned to the corresponding residues in S1. Next we obtained E2 from PDBID 5FER (S4) and aligned it to S1. All redundant structures were deleted, keeping only: DDB1 and Cul4A from S2, NEDD8 from S3, Rbx1 and Ubq from S1, and E2 from S4. Next, to account for its flexibility, we obtained 12 DDB1 structures as described by Bai et. al. [27] (3I8E, 3E0C, 3EI4, 4E54, 4TZ4, 6FCV, 4A08, 4A0B, 4A0L, 3EI3, 6PAI, 2B5L). All structures were aligned to S2 using BPB domain residues (395 to 705). We obtained the CRBN and DDB1 complex from PDBID 4TZ4 (S5). S2 was then aligned to each DDB1 conformation using BPA and BPC domains (2 to 394 and 706 to 1145). The model for each DDB1 structure was saved to a separate file. All the models generated are included in P4ward for the accessible lysine check.

## 2 Supplementary Tables

**Table S1:**
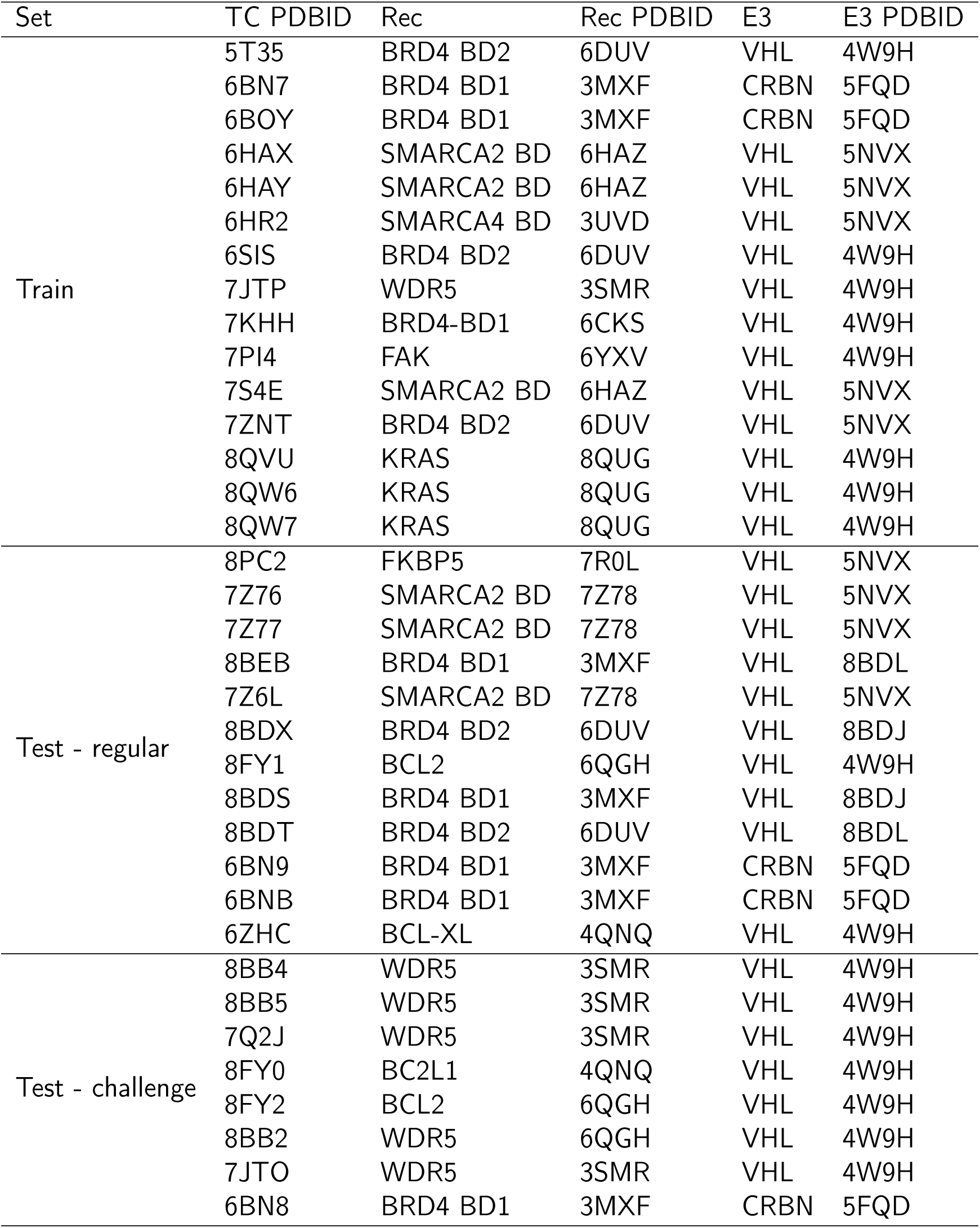
The PDB accession codes and description for all structures used in this study.

**Table S2:**
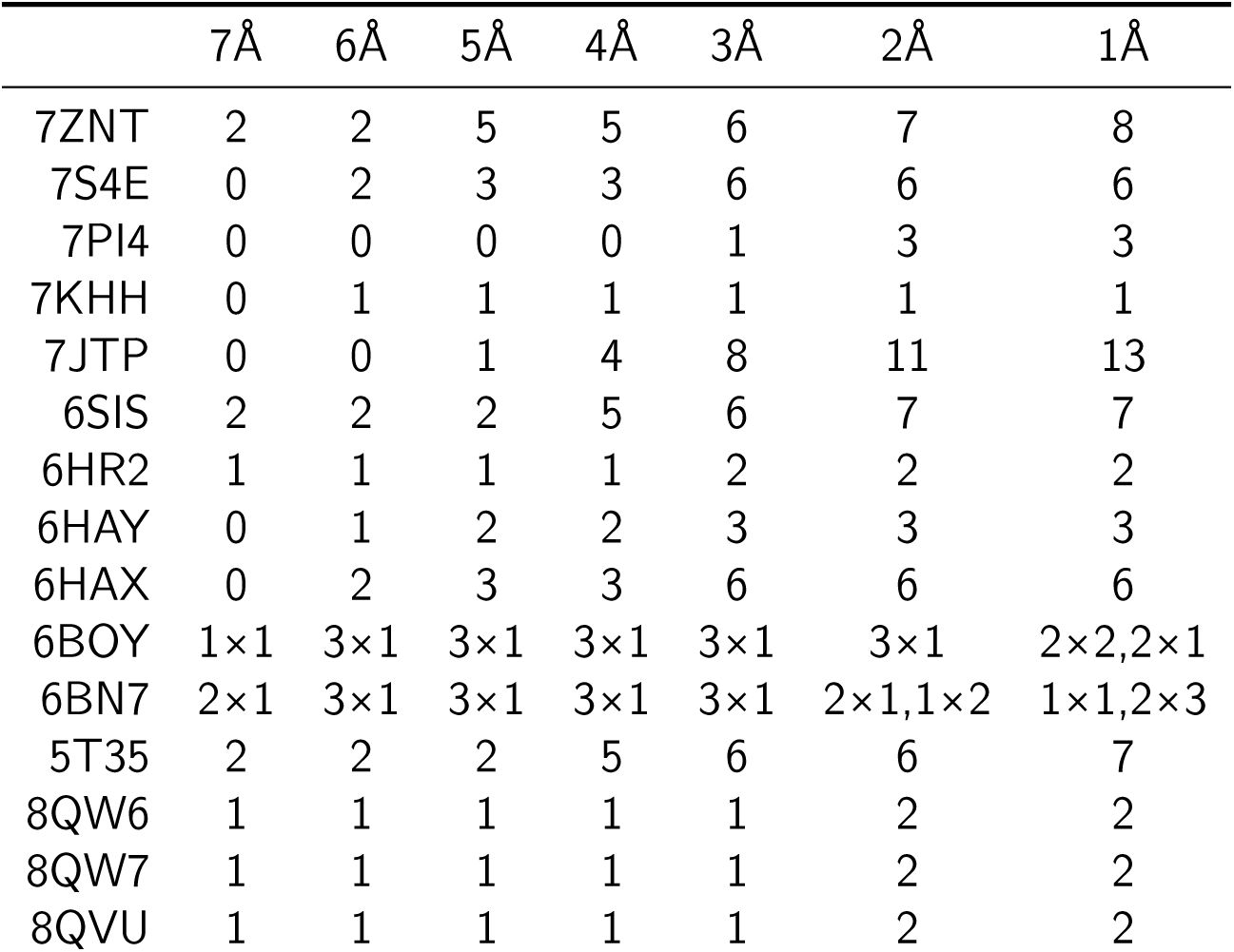
The number of accessible lysines identified for each RefE3_p_ in our benchmark set, for increasingly higher lysine occlusion cutoff values (Å). The notation for 6BOY and 6BN7 denotes (number of CRL models)×(number of accessible lysines), *i.e.* how many models had a specific number of accessible lysines.

**Table S3:**
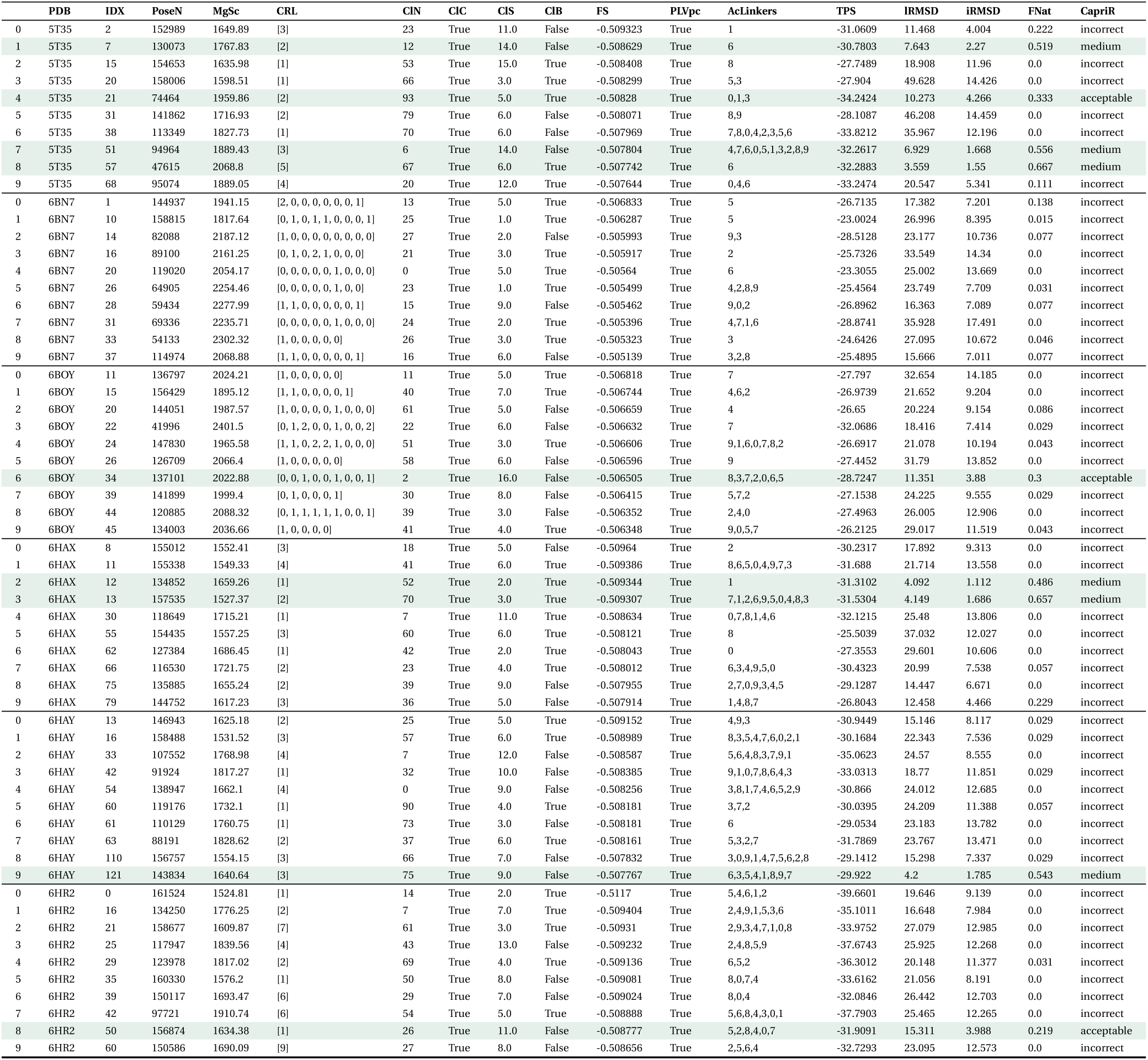
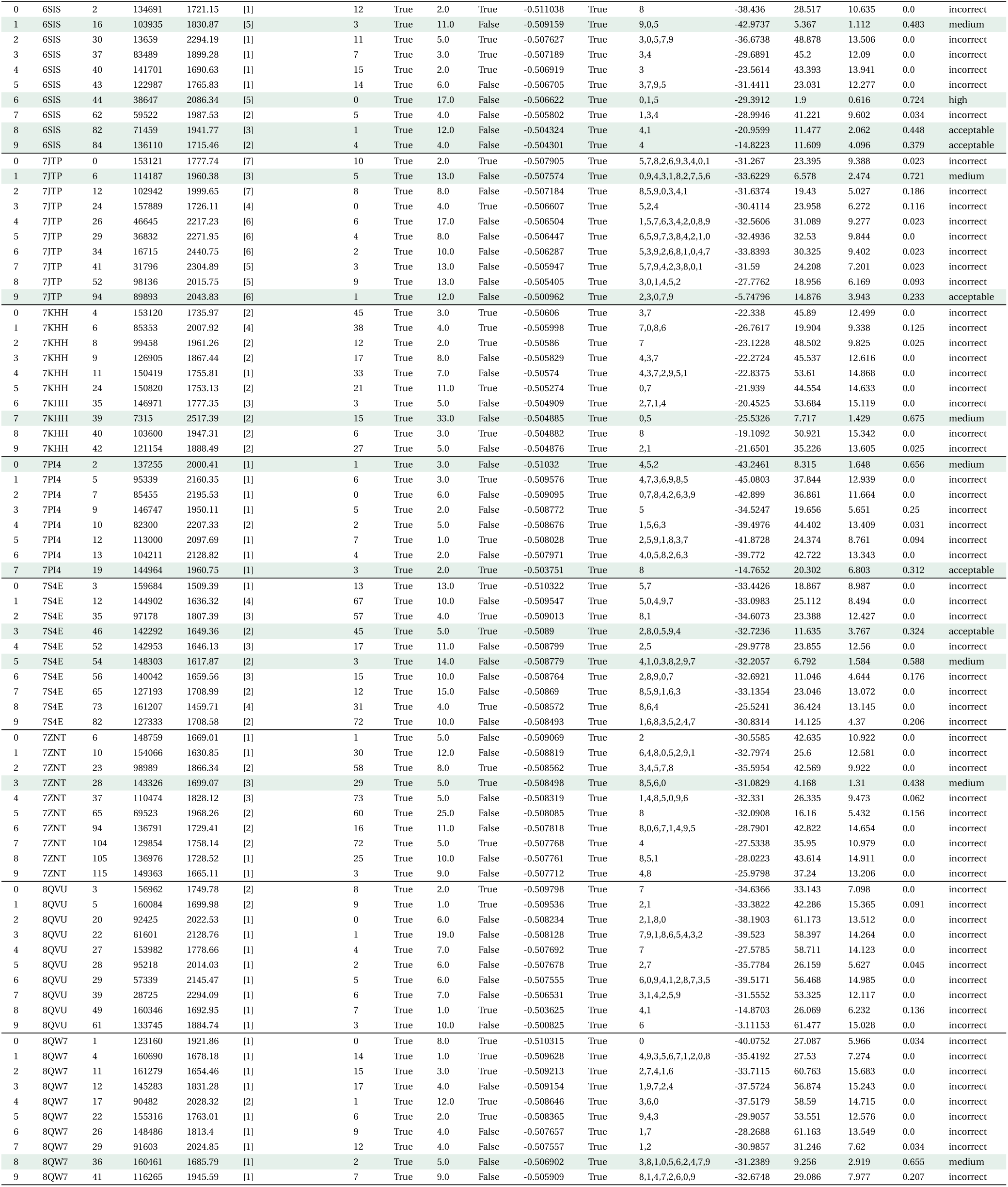
Full summary of results for configuration code 105 in the training set. For each reference PDB, we show the full result data written by P4ward for the top 10 predictions. **IDX**: the index for each pose without filtering by cluster centroids, ranked by the final score. **PoseN**: Megadock protein pose number. **MgSc**: Megadock score for the protein pose. Higher values correspond to more favourable poses. **CRL**: The number of accessible lysines found for each CRL model. **ClN**: The cluster number the pose belongs to, generated by trend clustering. **ClC**: If the protein pose is a cluster centroid. The final results include only the trend cluster centroids. **ClS**: The size of the cluster the protein pose belongs to. **ClB**: If the protein pose has the best final score in its cluster. **FS**: P4ward’s final score. Used to rank the results. **PLVpc**: If it was possible to sample at least one PLV_pc_ for the pose. **AcLinkers**: Out of the 10 PLV_pc_s sampled, which ones were sampled successfully. The numbers are ordered from best-scoring to worst-scoring PLV_pc_. **TPS**: Top PLV_pc_ score. **lRMSD**: ligand RMSD, from capri benchmark calculation. This corresponds to the RMSD of the smallest protein. **iRMSD**: interface RMSD, from capri benchmark calculation. **FNat**: Fraction of native contacts, from capri benchmark calculation. **CapriR**: Capri rank.

**Table S4:** Benchmarking of the top 10 TC predictions for PDBs 6HAX (Protac2), 6HAY (Protac1.) and 7S4E (ACBI1) against the three conformational clusters for each TC as predicted by Dixon *et al.* [22]. **ClusterN**: Cluster used as benchmarking reference. **PoseN**: Megadock pose number. **FNat**: Fraction of native contacts, from capri benchmark calculation. **lRMSD**: ligand RMSD, from capri benchmark calculation. This corresponds to the RMSD of the smallest protein. **iRMSD**: interface RMSD, from capri benchmark calculation. **Rank**: Capri rank. **CrysRank**: Previous capri rank, calculated using the crystal pose as reference.

|  | PDB | ClusterN | PoseN | FNat | IRMSD | iRMSD | Rank | CrystRank |
| --- | --- | --- | --- | --- | --- | --- | --- | --- |
| 0 | 6HAX | 0 | 155012 | 0.0 | 21.24 | 9.63 | incorrect | incorrect |
| 1 | 6HAX | 0 | 155338 | 0.0 | 21.87 | 15.625 | incorrect | incorrect |
| 2 | 6HAX | 0 | 134852 | 0.154 | 12.417 | 3.336 | acceptable | medium |
| 3 | 6HAX | 0 | 157535 | 0.173 | 12.757 | 4.425 | incorrect | medium |
| 4 | 6HAX | 0 | 118649 | 0.0 | 28.172 | 14.745 | incorrect | incorrect |
| 5 | 6HAX | 0 | 154435 | 0.019 | 30.537 | 13.576 | incorrect | incorrect |
| 6 | 6HAX | 0 | 127384 | 0.019 | 24.444 | 12.415 | incorrect | incorrect |
| 7 | 6HAX | 0 | 116530 | 0.0 | 30.865 | 10.58 | incorrect | incorrect |
| 8 | 6HAX | 0 | 135885 | 0.019 | 12.949 | 7.631 | incorrect | incorrect |
| 9 | 6HAX | 0 | 144752 | 0.327 | 5.968 | 3.191 | acceptable | incorrect |
| 10 | 6HAX | 1 | 155012 | 0.0 | 18.686 | 9.739 | incorrect | incorrect |
| 11 | 6HAX | 1 | 155338 | 0.0 | 23.049 | 13.621 | incorrect | incorrect |
| 12 | 6HAX | 1 | 134852 | 0.514 | 7.149 | 1.543 | medium | medium |
| 13 | 6HAX | 1 | 157535 | 0.486 | 7.594 | 2.379 | acceptable | medium |
| 14 | 6HAX | 1 | 118649 | 0.0 | 26.035 | 14.248 | incorrect | incorrect |
| 15 | 6HAX | 1 | 154435 | 0.0 | 38.685 | 11.852 | incorrect | incorrect |
| 16 | 6HAX | 1 | 127384 | 0.0 | 31.069 | 10.378 | incorrect | incorrect |
| 17 | 6HAX | 1 | 116530 | 0.086 | 17.85 | 6.545 | incorrect | incorrect |
| 18 | 6HAX | 1 | 135885 | 0.0 | 17.908 | 7.121 | incorrect | incorrect |
| 19 | 6HAX | 1 | 144752 | 0.086 | 16.25 | 5.057 | incorrect | incorrect |
| 20 | 6HAX | 2 | 155012 | 0.0 | 22.478 | 9.299 | incorrect | incorrect |
| 21 | 6HAX | 2 | 155338 | 0.0 | 20.498 | 14.098 | incorrect | incorrect |
| 22 | 6HAX | 2 | 134852 | 0.457 | 11.026 | 2.01 | acceptable | medium |
| 23 | 6HAX | 2 | 157535 | 0.543 | 10.591 | 2.41 | medium | medium |
| 24 | 6HAX | 2 | 118649 | 0.0 | 29.712 | 13.89 | incorrect | incorrect |
| 25 | 6HAX | 2 | 154435 | 0.0 | 31.497 | 11.4 | incorrect | incorrect |
| 26 | 6HAX | 2 | 127384 | 0.0 | 24.634 | 10.521 | incorrect | incorrect |
| 27 | 6HAX | 2 | 116530 | 0.0 | 28.488 | 8.17 | incorrect | incorrect |
| 28 | 6HAX | 2 | 135885 | 0.057 | 14.92 | 6.731 | incorrect | incorrect |
| 29 | 6HAX | 2 | 144752 | 0.4 | 7.502 | 3.554 | acceptable | incorrect |
| 30 | 6HAY | 0 | 146943 | 0.054 | 14.372 | 8.162 | incorrect | incorrect |
| 31 | 6HAY | 0 | 158488 | 0.054 | 17.366 | 7.198 | incorrect | incorrect |
| 32 | 6HAY | 0 | 107552 | 0.018 | 29.169 | 8.602 | incorrect | incorrect |
| 33 | 6HAY | 0 | 91924 | 0.018 | 22.152 | 13.218 | incorrect | incorrect |
| 34 | 6HAY | 0 | 138947 | 0.0 | 24.557 | 14.764 | incorrect | incorrect |
| 35 | 6HAY | 0 | 119176 | 0.089 | 21.655 | 12.845 | incorrect | incorrect |
| 36 | 6HAY | 0 | 110129 | 0.0 | 22.594 | 15.047 | incorrect | incorrect |
| 37 | 6HAY | 0 | 88191 | 0.0 | 27.952 | 15.132 | incorrect | incorrect |
| 38 | 6HAY | 0 | 156757 | 0.036 | 13.038 | 7.121 | incorrect | incorrect |
| 39 | 6HAY | 0 | 143834 | 0.25 | 14.454 | 4.544 | incorrect | medium |
| 40 | 6HAY | 1 | 146943 | 0.0 | 16.879 | 8.017 | incorrect | incorrect |
| 41 | 6HAY | 1 | 158488 | 0.0 | 25.011 | 7.437 | incorrect | incorrect |
| 42 | 6HAY | 1 | 107552 | 0.0 | 22.892 | 8.366 | incorrect | incorrect |
| 43 | 6HAY | 1 | 91924 | 0.029 | 19.082 | 12.085 | incorrect | incorrect |
| 44 | 6HAY | 1 | 138947 | 0.0 | 24.799 | 12.757 | incorrect | incorrect |
| 45 | 6HAY | 1 | 119176 | 0.029 | 26.092 | 11.617 | incorrect | incorrect |
| 46 | 6HAY | 1 | 110129 | 0.0 | 24.601 | 13.944 | incorrect | incorrect |
| 47 | 6HAY | 1 | 88191 | 0.0 | 23.152 | 13.573 | incorrect | incorrect |
| 48 | 6HAY | 1 | 156757 | 0.0 | 17.633 | 7.232 | incorrect | incorrect |
| 49 | 6HAY | 1 | 143834 | 0.543 | 3.848 | 1.76 | medium | medium |
| 50 | 6HAY | 2 | 146943 | 0.049 | 14.223 | 8.297 | incorrect | incorrect |
| 51 | 6HAY | 2 | 158488 | 0.073 | 20.958 | 7.625 | incorrect | incorrect |
| 52 | 6HAY | 2 | 107552 | 0.024 | 26.95 | 8.236 | incorrect | incorrect |
| 53 | 6HAY | 2 | 91924 | 0.024 | 21.273 | 13.168 | incorrect | incorrect |
| 54 | 6HAY | 2 | 138947 | 0.0 | 26.203 | 14.63 | incorrect | incorrect |
| 55 | 6HAY | 2 | 119176 | 0.073 | 23.583 | 12.778 | incorrect | incorrect |
| 56 | 6HAY | 2 | 110129 | 0.0 | 21.66 | 14.951 | incorrect | incorrect |
| 57 | 6HAY | 2 | 88191 | 0.0 | 25.812 | 15.022 | incorrect | incorrect |
| 58 | 6HAY | 2 | 156757 | 0.049 | 15.231 | 7.379 | incorrect | incorrect |
| 59 | 6HAY | 2 | 143834 | 0.244 | 14.001 | 4.612 | incorrect | medium |
| 60 | 7S4E | 0 | 159684 | 0.0 | 21.697 | 9.863 | incorrect | incorrect |
| 61 | 7S4E | 0 | 144902 | 0.0 | 22.917 | 9.07 | incorrect | incorrect |
| 62 | 7S4E | 0 | 97178 | 0.0 | 25.223 | 11.745 | incorrect | incorrect |
| 63 | 7S4E | 0 | 142292 | 0.375 | 15.056 | 3.33 | acceptable | acceptable |
| 64 | 7S4E | 0 | 142953 | 0.0 | 26.009 | 11.962 | incorrect | incorrect |
| 65 | 7S4E | 0 | 148303 | 0.375 | 8.313 | 2.41 | acceptable | medium |
| 66 | 7S4E | 0 | 140042 | 0.031 | 15.214 | 5.666 | incorrect | incorrect |
| 67 | 7S4E | 0 | 127193 | 0.0 | 21.938 | 13.271 | incorrect | incorrect |
| 68 | 7S4E | 0 | 161207 | 0.0 | 39.059 | 12.688 | incorrect | incorrect |
| 69 | 7S4E | 0 | 127333 | 0.406 | 9.62 | 3.357 | acceptable | incorrect |
| 70 | 7S4E | 1 | 159684 | 0.0 | 17.004 | 9.506 | incorrect | incorrect |
| 71 | 7S4E | 1 | 144902 | 0.016 | 30.906 | 8.95 | incorrect | incorrect |
| 72 | 7S4E | 1 | 97178 | 0.0 | 23.636 | 14.864 | incorrect | incorrect |
| 73 | 7S4E | 1 | 142292 | 0.246 | 11.84 | 4.655 | incorrect | acceptable |
| 74 | 7S4E | 1 | 142953 | 0.016 | 23.013 | 14.826 | incorrect | incorrect |
| 75 | 7S4E | 1 | 148303 | 0.164 | 15.184 | 4.777 | incorrect | medium |
| 76 | 7S4E | 1 | 140042 | 0.197 | 10.133 | 4.694 | incorrect | incorrect |
| 77 | 7S4E | 1 | 127193 | 0.0 | 27.348 | 15.469 | incorrect | incorrect |
| 78 | 7S4E | 1 | 161207 | 0.016 | 31.141 | 14.29 | incorrect | incorrect |
| 79 | 7S4E | 1 | 127333 | 0.066 | 24.541 | 7.474 | incorrect | incorrect |
| 80 | 7S4E | 2 | 159684 | 0.049 | 18.586 | 7.964 | incorrect | incorrect |
| 81 | 7S4E | 2 | 144902 | 0.024 | 28.833 | 8.34 | incorrect | incorrect |
| 82 | 7S4E | 2 | 97178 | 0.0 | 21.132 | 12.147 | incorrect | incorrect |
| 83 | 7S4E | 2 | 142292 | 0.146 | 10.308 | 5.083 | incorrect | acceptable |
| 84 | 7S4E | 2 | 142953 | 0.0 | 21.232 | 12.287 | incorrect | incorrect |
| 85 | 7S4E | 2 | 148303 | 0.439 | 11.501 | 2.051 | acceptable | medium |
| 86 | 7S4E | 2 | 140042 | 0.317 | 10.799 | 3.807 | acceptable | incorrect |
| 87 | 7S4E | 2 | 127193 | 0.0 | 25.879 | 12.895 | incorrect | incorrect |
| 88 | 7S4E | 2 | 161207 | 0.0 | 32.728 | 12.806 | incorrect | incorrect |
| 89 | 7S4E | 2 | 127333 | 0.146 | 20.166 | 5.668 | incorrect | incorrect |

**Table S5:**
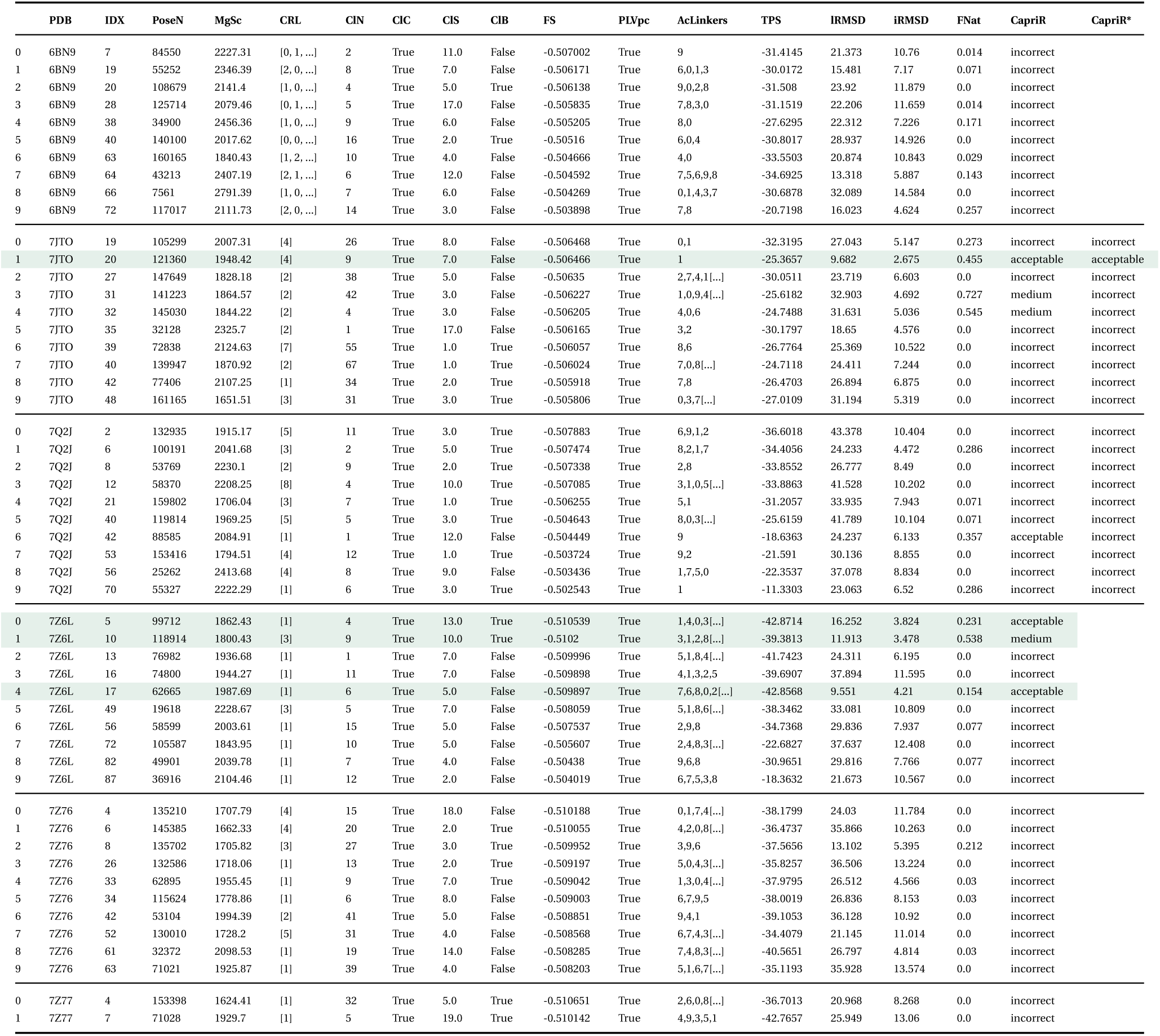
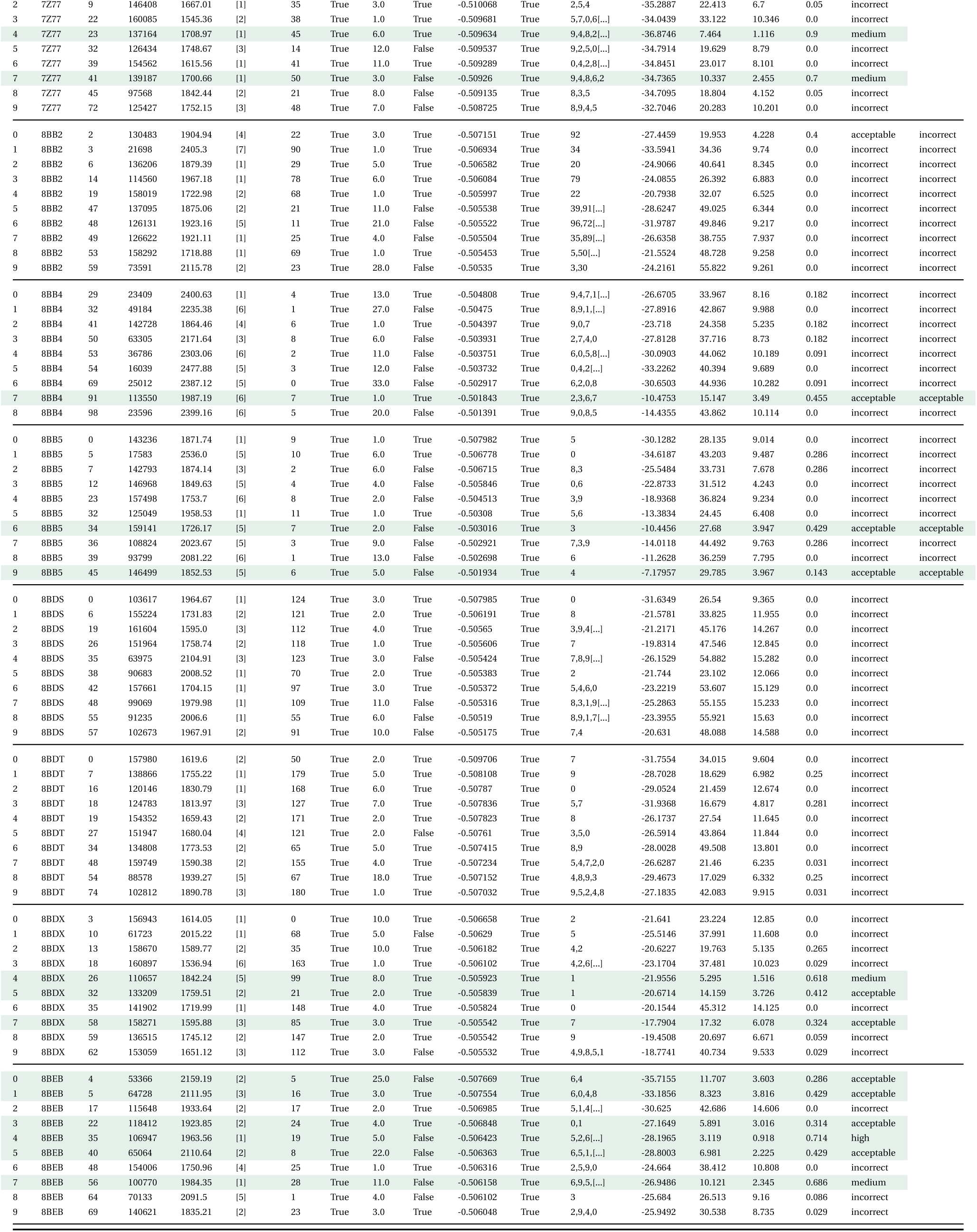
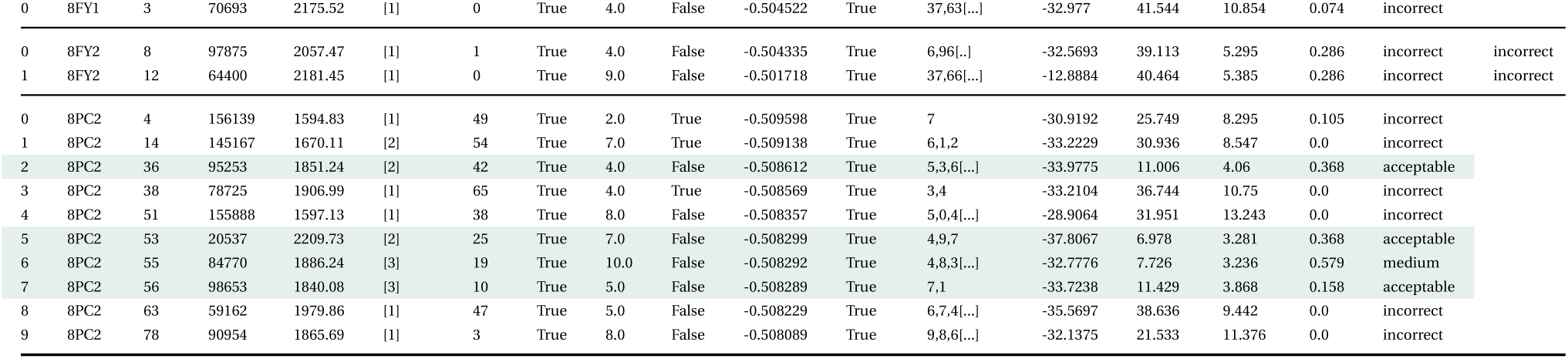
Full summary of results for configuration code 105 on the benchmark test set. For each reference PDB, we show the full result data written by P4ward for the top 10 predictions. **IDX**: the index for each pose without filtering by cluster centroids, ranked by the final score. **PoseN**: Megadock protein pose number. **MgSc**: Megadock score for the protein pose. Higher values correspond to more favourable poses. **CRL**: The number of accessible lysines found for each CRL model. **ClN**: The cluster number the pose belongs to, generated by trend clustering. **ClC**: If the protein pose is a cluster centroid. The final results include only the trend cluster centroids. **ClS**: The size of the cluster the protein pose belongs to. **ClB**: If the protein pose has the best final score in its cluster. **FS**: P4ward’s final score. Used to rank the results. **PLVpc**: If it was possible to sample at least one PLV_pc_ for the pose. **AcLinkers**: Out of the 10 PLV_pc_s sampled, which ones were sampled successfully. The numbers are ordered from best-scoring to worst-scoring PLV_pc_. **TPS**: Top PLV_pc_ score. **lRMSD**: ligand RMSD, from capri benchmark calculation. This corresponds to the RMSD of the smallest protein. **iRMSD**: interface RMSD, from capri benchmark calculation. **FNat**: Fraction of native contacts, from capri benchmark calculation. **CapriR**: Capri rank. **CapriR\***: Capri rank without the FNat contribution. Please see text section **??** for details. The columns **CRL** and **AcLinkers** were truncated to preserve space.

**Table S6:** Comparison between the benchmarking studies performed by Rovers and Schapira [25] and the benchmark of P4ward in this study.

| TC PDBID | Ref. Benchmark | P4ward's Benchmark | Note |
| --- | --- | --- | --- |
| 5T35 | ✓ | ✓ |  |
| 6BN7 | ✓ | ✓ |  |
| 6BN8 | ✓ |  | intractable P4ward run time |
| 6BN9 | ✓ | ✓ |  |
| 6BNB | ✓ | ✓ |  |
| 6BOY | ✓ | ✓ |  |
| 6HAX | ✓ | ✓ |  |
| 6HAY | ✓ | ✓ |  |
| 6HR2 | ✓ | ✓ |  |
| 6SIS | ✓ | ✓ |  |
| 6W7O | ✓ |  | not vhl/crbn |
| 6W8I | ✓ |  | not vhl/crbn |
| 6ZHC | ✓ | ✓ |  |
| 7JTO | ✓ | ✓ |  |
| 7JTP | ✓ | ✓ |  |
| 7KHH | ✓ | ✓ |  |
| 7PI4 | ✓ | ✓ |  |
| 7Q2J | ✓ | ✓ |  |
| 7S4E | ✓ | ✓ |  |
| 7Z6L | ✓ | ✓ |  |
| 7Z76 | ✓ | ✓ |  |
| 7Z77 | ✓ | ✓ |  |
| 7ZNT | ✓ | ✓ |  |
| 8BB2 | ✓ | ✓ |  |
| 8BB3 | ✓ |  | isomer of 8BB2 |
| 8BB4 | ✓ | ✓ |  |
| 8BB5 | ✓ | ✓ |  |
| 8BDS | ✓ | ✓ |  |
| 8BDT | ✓ | ✓ |  |
| 8BDX | ✓ | ✓ |  |
| 8BEB | ✓ | ✓ |  |
| 8DSO | ✓ |  | not vhl/crbn |
| 8FY0 | ✓ | ✓ |  |
| 8FY1 | ✓ | ✓ |  |
| 8FY2 | ✓ | ✓ |  |
| 8PC2 | ✓ | ✓ |  |
| 8QJR | ✓ |  | no interface |
| 8QU8 | ✓ |  | CRL complex of 8QVU |
| 8QVU | ✓ | ✓ |  |
| 8QW6 | ✓ |  | no linker sampling possible |
| 8QW7 | ✓ | ✓ |  |
| 8R5H | ✓ |  | CRL complex of 5T35 |
| 8RWZ | ✓ |  | CRL complex of 5T35 |

## 3 Supplementary Figures

**Figure S1:**
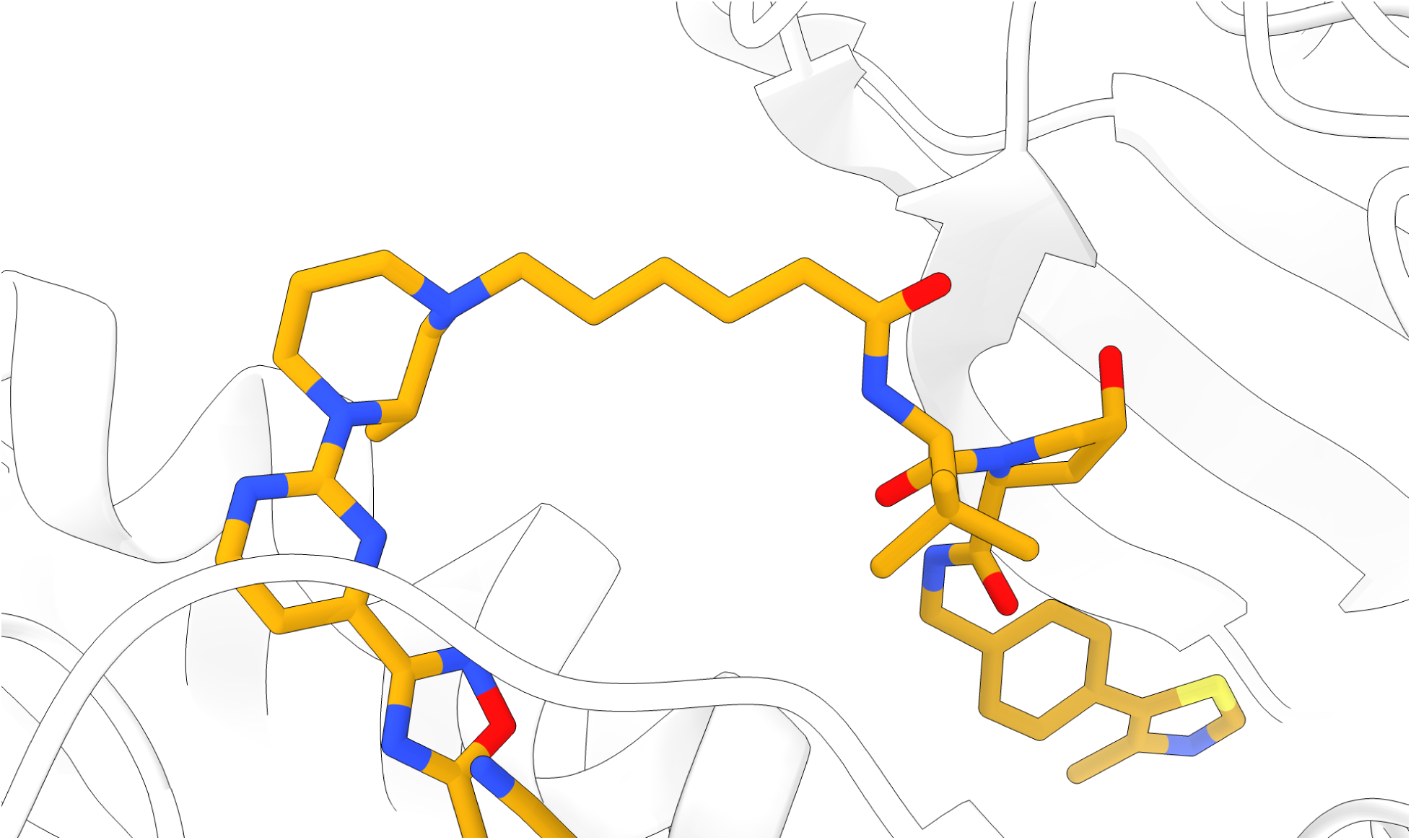
Bound protac structure (orange) from in PDBID 8QW6. This linker’s stretched conformation cannot be reproduced when sampling with P4ward.

**Figure S2:**
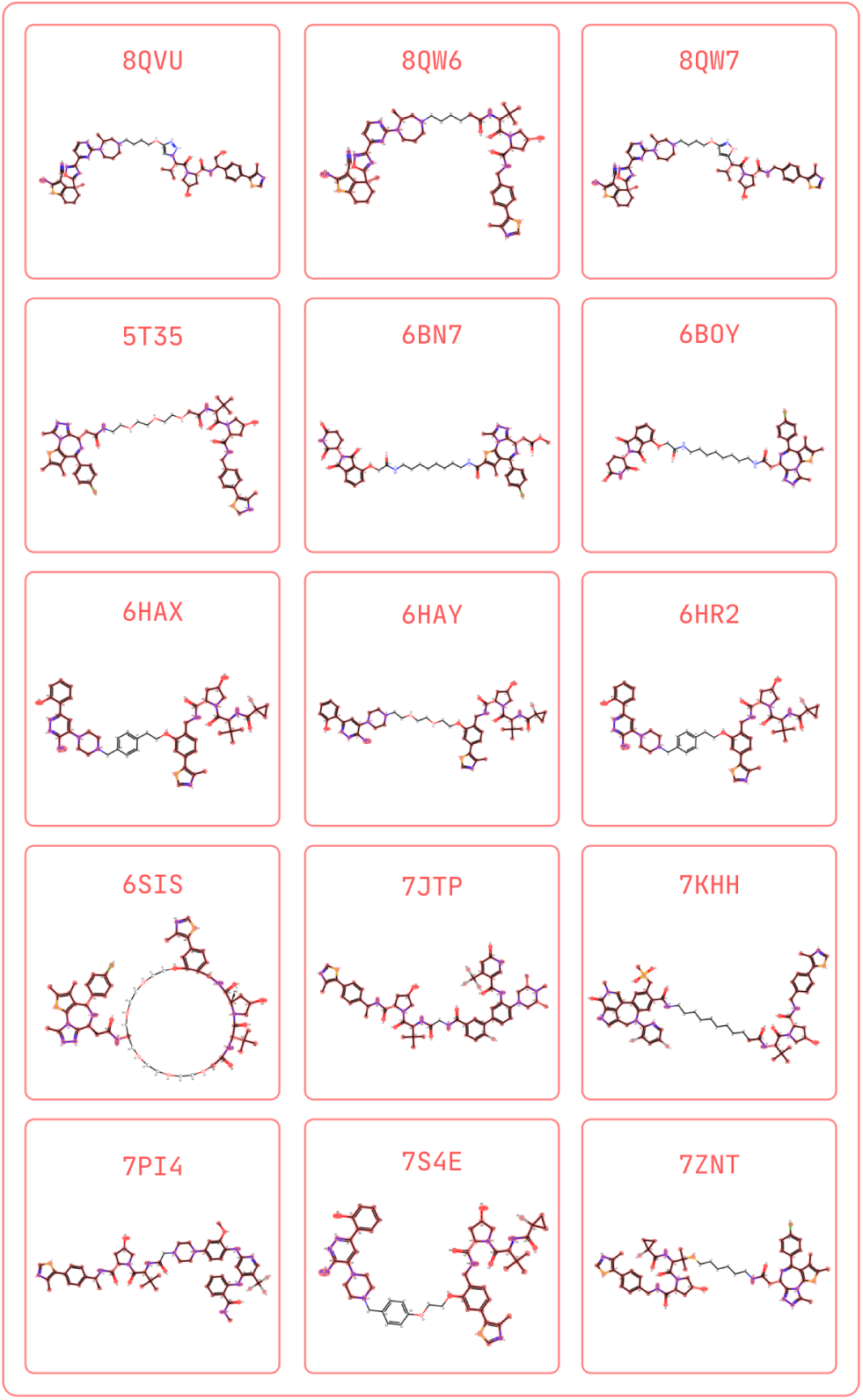
Using RDKit, P4ward accurately matches the ligands’ atoms onto the full protac structure. The matched atoms and bonds are represented in red. In subsequent sampling steps, the non-matched atoms will be considered part of the linker and therefore treated as flexible for sampling, while the matched atoms are considered part of either ligand and will maintain the coordinates from the input .mol2 files.

**Figure S3:**
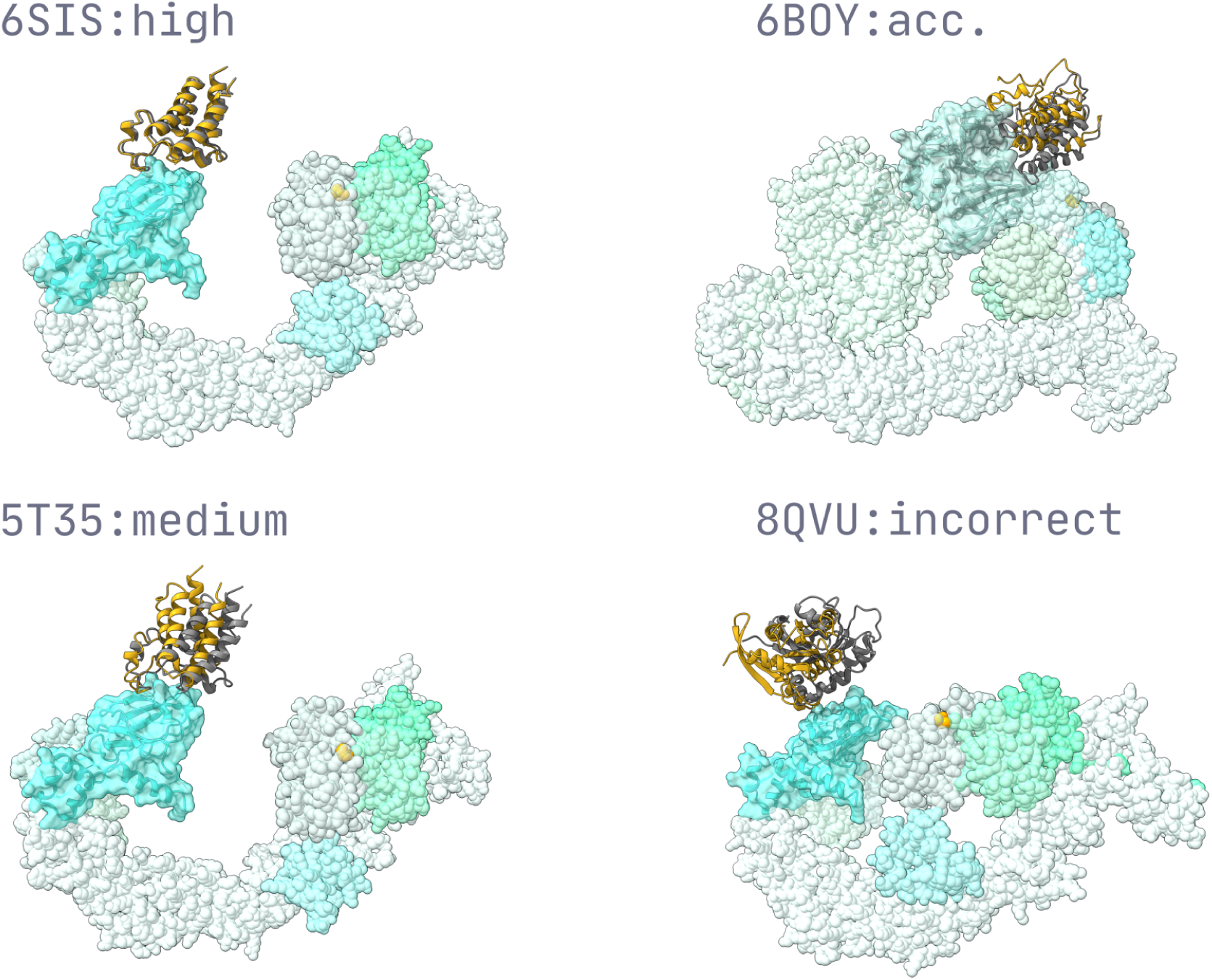
Full CRL complex models for each example prediction from conf. 105 shown in Figure 4.

**Figure S4:**
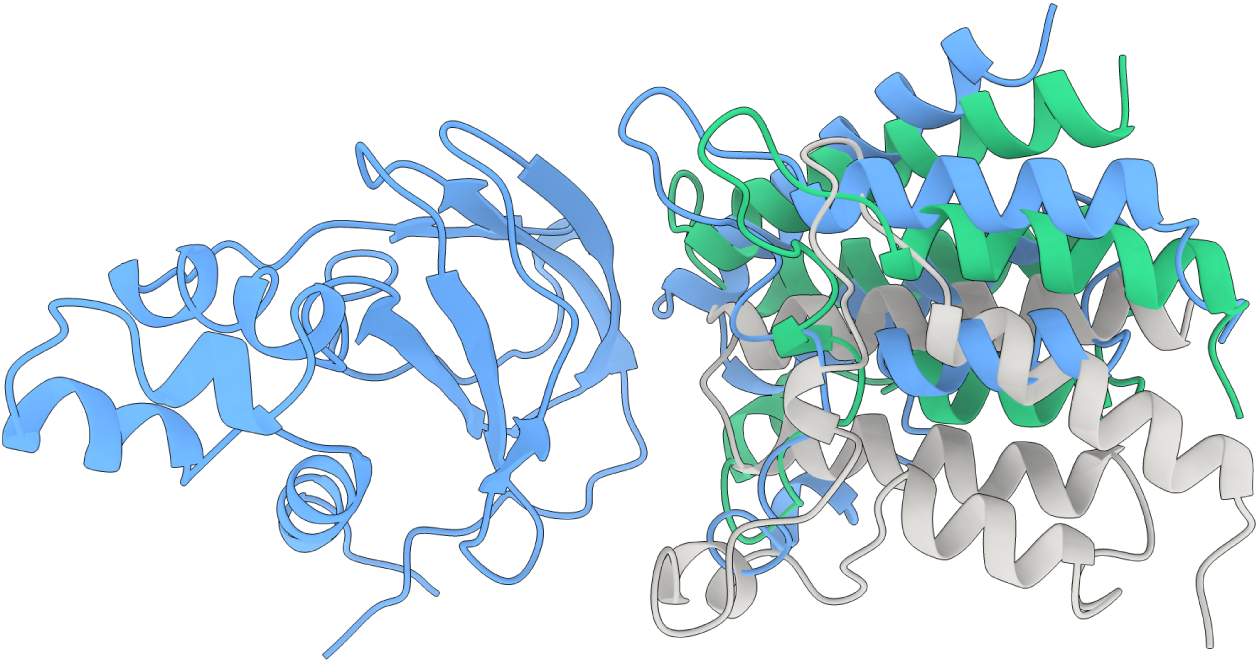
Comparison between Dixon *et al.*’s cluster 0 for Protac 2 (blue), The reference crystal pose (6HAX, gray), and P4ward conformation at rank 9 (green).

**Figure S5:**
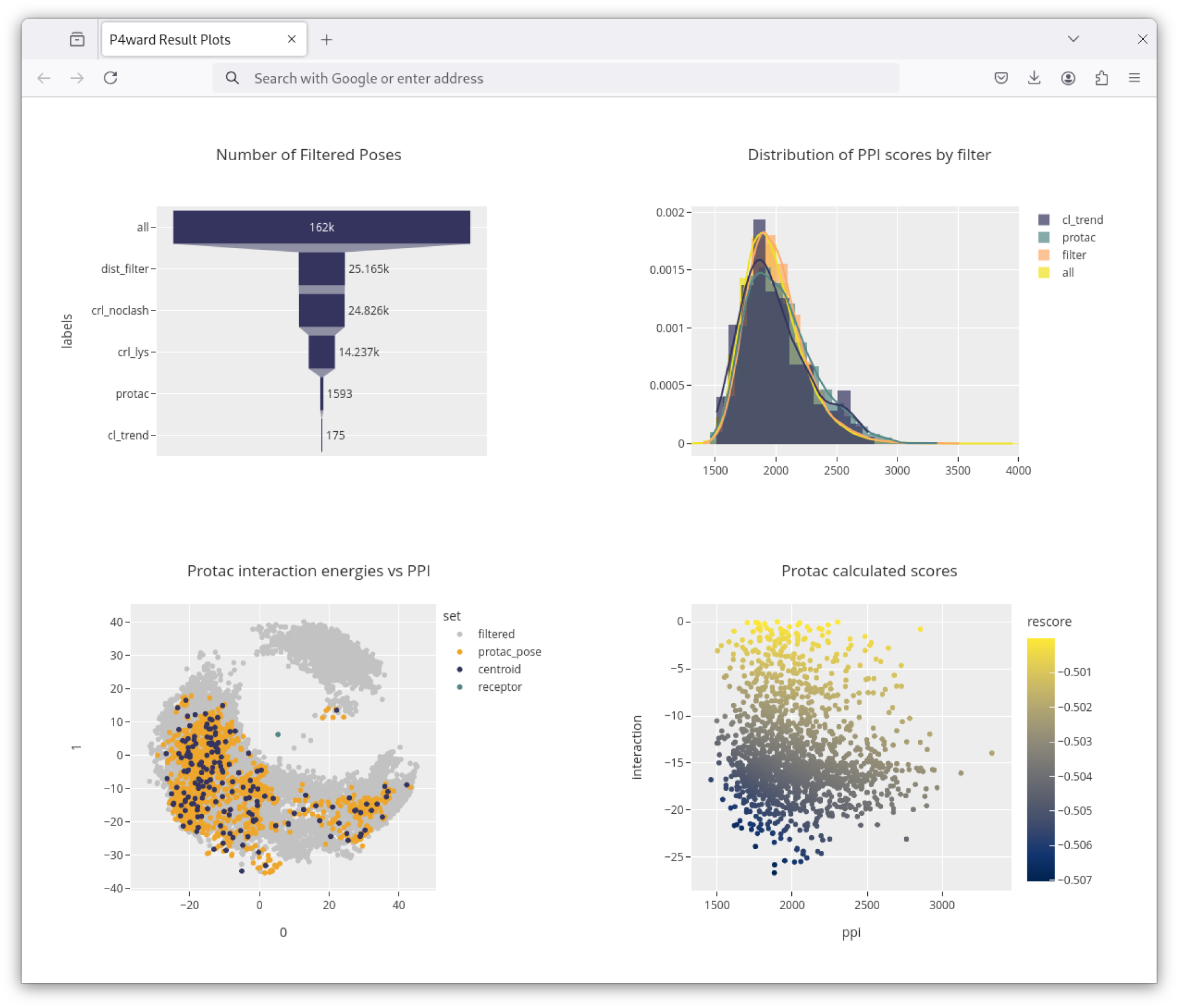
Example of interactive output generated automatically by P4ward (for reference 8BDX). This output consist of a single file which is opened in the web browser. (Top left) The number of protein poses remaining after each pipeline stage. (Top right) The distributions of the Megadock score for the protein poses in the main pipeline stages. (Bottom left) Principal components 1 and 2 from principal component analysis of the 3D coordinates of the protein poses in the main pipeline stages. (Bottom right) Scatter plot of the scores in the final predicted models: protein-protein score, protein-protac interaction score, and rescore by final score.

## Acronyms

bE3: bound E3 structure. 18, 22
bE3L: bound E3 structure’s ligand. 22
bRec: bound Rec structure. 18, 22
bRecL: bound Rec structure’s ligand. 22
E3: E3 Ligase Substrate Receptor. 17–20, 22, 23
E3_p_: E3 docked pose. 18–22
E3L: E3 Ligand. 17–19, 22
E3L_p_: E3 Ligand pose. 19, 20, 22
PLV: Protac Linker Variant. 17–22
PLV_pc_: Protac Linker Variant pose’s linker conformation. 18, 20, 21
PLV_p_: Protac Linker Variant pose. 18, 20, 21
Rec: Receptor. 17–23
Rec_pCRL_: Receptor bound to E3_p_ aligned to the CRL model. 20
RecL: Receptor Ligand. 17–20
RefE3_p_: Reference E3 binding mode. 18, 22
RefE3L_p_: Reference E3 binding mode’s ligand. 22
uE3: unbound E3 structure. 18, 23 **uE3L** unbound E3 structure’s ligand. 23
uRec: unbound Rec structure. 18, 23
uRecL: unbound Rec structure’s ligand. 23

## Notes

### Competing Interest Statement

The authors have declared no competing interest.

